# iG6PSnFR: A genetically encoded fluorescent sensor for observing glucose-6-phosphate dynamics in living preparations

**DOI:** 10.64898/2026.07.22.740099

**Authors:** Jonathan S. Marvin, Anastasia Tsives, Shih Ming Huang, Zhanat Koshenov, Raghabendra Adhikari, Aaron D. Wolfe, Ian J. Gonzalez, Daniel Feliciano, Timothy A. Ryan, Matthew J. Merrins, Daniel A. Colón-Ramos, Timothy A. Brown

**Affiliations:** Howard Hughes Medical Institute, Janelia Research Campus, Ashburn, VA, USA; Department of Neuroscience and Department of Cell Biology, Yale University School of Medicine; New Haven, CT, USA; Department of Cellular & Molecular Physiology, Yale School of Medicine, New Haven, CT, USA; Department of Biochemistry & Biophysics, Weill Cornell Medicine, New York, NY, USA

## Abstract

Glucose-6-phosphate (G6P) is a key intermediate in multiple energetic and anabolic pathways, and quantifying its dynamics is essential for understanding cellular physiology. We have previously developed a syndicate of intensity-based, genetically encoded sensors based on the insertion of circularly permuted GFP into a Venus-flytrap-like analyte-binding protein. Here we use the same approach to develop an intensity-based G6P Sensing Fluorescent Reporter (iG6PSnFR). We present two variants: a G6P-activated sensor that increases fluorescence and a G6P-inactivated sensor that decreases fluorescence. We validate performance across progressively more complex preparations, including purified protein in vitro, immortalized and primary neuronal cultures, isolated pancreatic islets, in an intravital liver model, and finally in vivo in *C. elegans* neurons. In each context, iG6PSnFR reports G6P changes consistent with expected responses to physiological perturbations.

## Introduction

Glucose-6-phosphate (G6P) occupies a central role in cellular metabolism and is a critical intermediate in glycolysis, gluconeogenesis, and the pentose phosphate and hexosamine pathways (*1*). Its regulation is vital for maintaining cellular energy homeostasis and overall metabolic function. Consequently, accurate measurement of intracellular G6P is essential for monitoring cellular responses to various biochemical perturbations in healthy and metabolically diseased tissues. Traditional methods for detecting G6P, including enzymatic assays and chromatography-based techniques, often involve complex sample preparation, long analysis times, and specialized instrumentation (*2*). These limitations make high-throughput or real-time monitoring of G6P challenging, particularly in biological systems where fast and non-invasive detection with cellular resolution is often desirable.

Genetically encoded fluorescence-based sensors are a promising alternative, offering high sensitivity, rapid response kinetics, and the ability to monitor metabolites in real-time both *in vitro* and *in vivo* contexts (*3*). Here we describe the design, optimization, and application of a fluorescence sensor for the detection of G6P. By fusing a G6P-specific binding protein with circularly permuted GFP, we created a pair of intensity-based G6P-Sensing Fluorescent Reporters (iG6PSnFRs). One is a traditional sensor that gets brighter when bound to G6P and the other is an inverse sensor, getting dimmer when bound to G6P. They can measure G6P levels in milliseconds, with potential applications including basic metabolic research, drug discovery, and potentially clinical diagnostics.

## Results

### Sensor development and in vitro characterization

We initially sought to develop a sensor for G6P by redesigning our existing sensor for glucose, iGlucoSnFR2 (*4*). To guide redesign of the ligand-binding pocket, we surveyed the protein structure database for proteins known to bind G6P. This search identified two candidates: a G6P-binding protein (HptA) from *Staphylococcus aureus* (*5*) and one from *Actinobacillus pleuropneumoniae* (AfuA) (*6*). Both are reported to be periplasmic binding proteins with Venus-flytrap mechanisms for binding their ligands, but HptA is reported to only bind G6P and galactose-6-phosphate, while AfuA is reported to also bind fructose-6-phosphate. As such, we went forward with plans to turn HptA (6LKK.pdb) into an intensity-based sensor.

Leveraging prior success with circularly-permuted GFP (cpGFP)-based sensors derived from Venus-flytrap like proteins (*4*, *7–9*), we chose four positions within HptA to insert cpSFGFP: after residues 93, 196, 292, or 298 (Fig. 1A, numbering based on 6LKK.pdb). We randomized residues at the junctions between the binding protein and cpSFGFP for each insertion site and screened approximately 200 variants for each site. Insertions at residues 93, 292, and 298 yielded functional sensors, whereas insertion at residue 196 did not (Supplementary Fig. S1A). Positions 292 and 298 are located at the interface of the two domains of the binding protein and are expected to allosterically couple ligand binding to changes in fluorescence when cpSFGFP is inserted. Sensors identified at both sites were selected for further optimization, culminating in a sensor with a large increase in fluorescence upon saturation with G6P (“iG6PSnFR”, ΔF/F ∼15, *K*_d_ ∼ 10 µM) and another with a large decrease in fluorescence upon saturation (“iG6PSnFR-D”, ΔF/F ∼ - 0.65, *K*_d_ ∼ 1 µM). Stopped-flow kinetic analysis shows that the sensors reach equilibrium within 100 msec. (Supplementary Fig. S1B). The excitation and emission profiles at both the GFP peak and with parameters typically used for YFP are reported in Supplementary Figs. S1C & S1D. Since some metabolic perturbations can affect pH, and GFPs including cpSFGFP are pH-sensitive, we have also reported the pH dependence of the sensors in Supplementary Fig. S1E. Both iG6PSnFR sensors exhibit a small change in their fluorescence lifetimes between the unbound and bound states, but the change is too small (∼0.1 nsec.) to function practically as a lifetime sensor (Supplementary Fig. S1F).

**Fig. 1.**
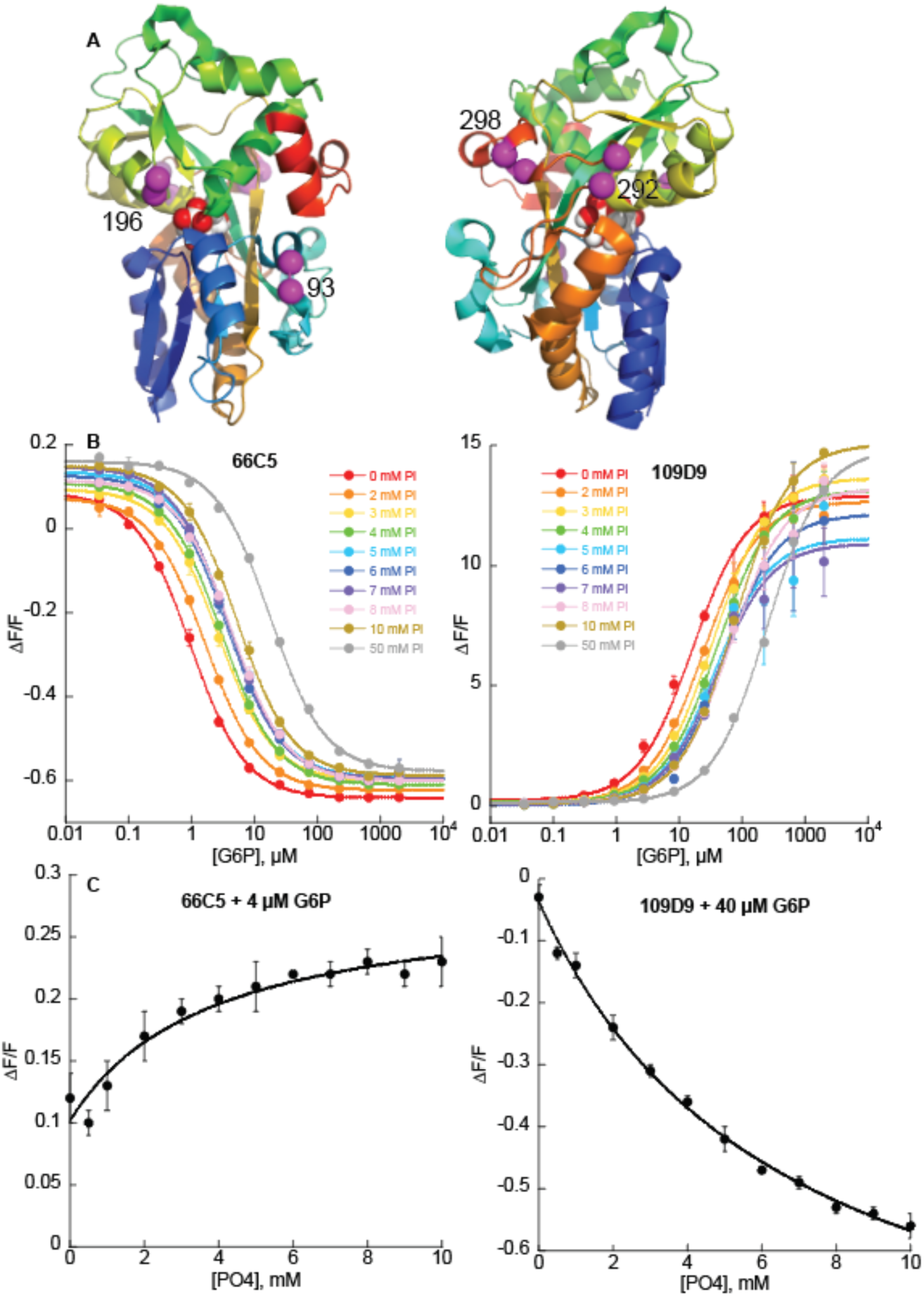
Design and *in vitro* characterization of iG6PSnFR. (**A**) Ribbon diagram of HptA (6LKK.pdb) with bound G6P shown as white and red spheres, and with numbered purple spheres indicating insertion points for cpSFGFP. (**B**) Titration of iG6PSnFR with G6P with varying concentrations of inorganic phosphate in TBS. Left is sensor iG6PSnFR-D, right is sensor iG6PSnFR. (**C**) Titration of iG6PSnFR with P_i_ in the presence of 4 µM or 40 µM G6P as indicated.

### Ligand binding specificity

Sensor specificity is a critical parameter for accurate metabolite detection. HptA was reported to be highly specific for G6P as determined by isothermal titration calorimetry (ITC) (*5*). However, ITC is limited in its ability to detect weak (ie, 100 micromolar to low millimolar) binding or competitive inhibition. Thus, we tested a panel of decoys relevant to metabolism: glucose, glucose-1-phosphate, fructose-6-phosphate, fructose-1,6-bisphosphate, and 6-phosphogluconate. We were unable to acquire galactose-6-phosphate, a ligand reported to bind to HptA (*5*). The sensors do not change fluorescence in response to any of these compounds (Supplementary Fig. S1G). They also do not bind the often-used inhibitor of glycolysis 2-deoxyglucose, nor its phosphorylated counterpart, 2-deoxyglucose-6-phoshate (Supplementary Fig. S1G). The sensor’s affinity for G6P is not affected by 10 mM glucose, the compound most likely to be present at high concentrations in cells (Supplementary Fig. S1H). We fused red fluorescent proteins (mRuby3 or mScarlet-I3), an infrared fluorescent protein (mIRFP670nano3), or HaloTag to the sensors’ C-terminus to provide optically inert options for ratiometry or normalization of expression and focus. The fusions’ effect on G6P affinity and ΔF/F is minimal (Supplementary Fig. S1I).

The affinity of both iG6PSnFRs for G6P decreases about 20-fold in the presence of 50 mM inorganic phosphate (P_i_), as shown in Fig. 1b. The *K*_d_ values change from 1 µM to 17 µM for iG6PSnFR-D and from 16 µM to 250 µM for iG6PSnFR. The effect of P_i_ on the sensors’ affinity for G6P means that a change in sensor fluorescence could result from micromolar changes in G6P concentration (as expected and desired) or from millimolar changes in P_i_ concentration. A recently published lifetime sensor for inorganic phosphate (Pi-Tq) indicates that under prolonged electrical stimulation, the concentration of P_i_ in cultured neurons increases from 4 mM to 6 mM (*10*). Thus, we titrated G6P while varying P_i_ from 0 mM to 50 mM, collecting more curves in the physiologically relevant concentration of P_i_ (Fig. 1B). Conversely, we titrated P_i_ with the concentration of G6P near the sensors’ *K*_d_ (Fig. 1C), which would be the point at which iG6PSnFR is most sensitive to fluctuations in P_i_. iG6PSnFR-D (with 4 µM G6P in solution) shows a minor 0.01 unit change in ΔF/F upon a shift from 4 mM to 6 mM P_i_, while iG6PSnFR (with 40 µM G6P in solution) shows a-0.11 unit change in ΔF/F.

### iG6PSnFRs performance in mammalian cell culture

To assess sensor performance in a cellular context, we expressed iG6PSnFR variants in immortalized cell culture. Both variants were fused to HaloTag to allow normalization of expression levels using a far-red fluorophore (JFX650) (*11*). In parallel, we also transfected HeLa cells with the glucose sensor, iGlucoSnFR2. We imaged these cells in a Cytation multi-well plate reader and collected both the green (to detect G6P or glucose) and red fluorescence to detect JFX650 coupled to HaloTag to normalize for variable protein expression. We then performed different perturbations that were expected to result in changes in cytosolic concentrations of G6P.

First, blocking glucose from entering cells by inhibition of the GLUT family of glucose transporters with the drug Glutor (*12*) results in rapid depletion of G6P as G6P stops being produced from glucose yet is still consumed by multiple pathways (Fig. 2A, blue curves). Inhibition of mitochondrial hexokinase by the drug lonidamine (*13*) also immediately reduced G6P levels (Fig. 2B, pink curves). Addition of 2-deoxyglucose (Fig. 2C, red curves) had a similar effect, but over a slightly longer time course, as the inhibition of hexokinase comes not by 2-deoxyglucose itself, but rather by the accumulation of 2-deoxyglucose-6-phosphate, the product of phosphorylation of 2-deoxyglucose (*14*). The difference in time course is more pronounced when the curves for each treatment are plotted on the same graph (Supplementary Fig. 2). Finally, equilibrating the cells in glucose-free buffer and then adding glucose (Fig. 2D, black lines) increases cytosolic G6P levels. Both sensors respond as expected, with iG6PSnFR-D showing inverse responses compared to iG6PSnFR.

**Fig. 2.**
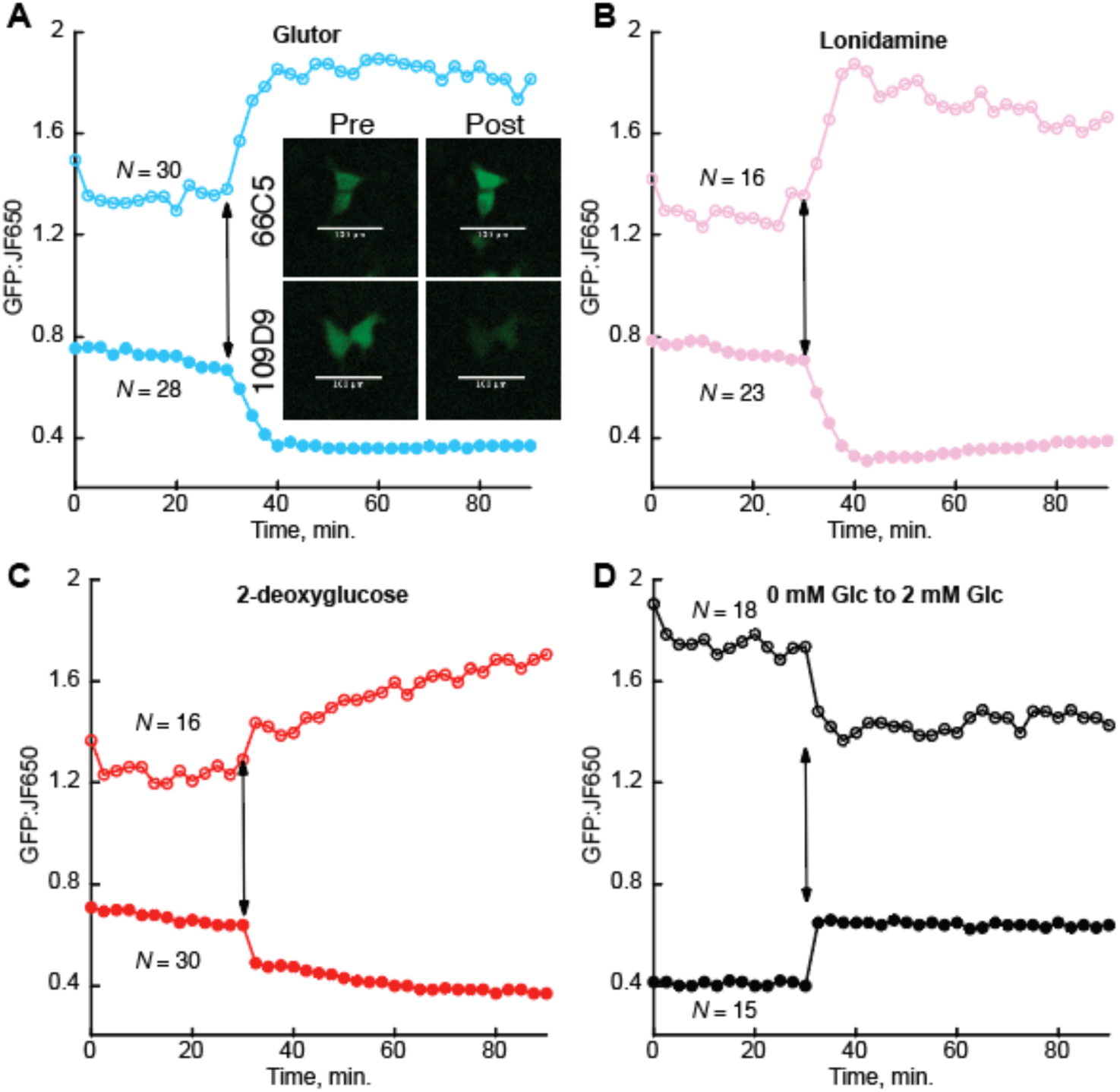
Effect of metabolic perturbations on G6P levels in HeLa cells. HeLa cells transfected with (cyto).iG6PSnFR.HaloTag or (cyto).iG6PSnFR-D.HaloTag and labelled with JFX650 were equilibrated with HEPES buffer containing 2 mM glucose and imaged in a Cytation multi-well plate imager. After 30 minutes (as indicated by the arrows), drugs were added to final concentrations of (**A**) 10 µM Glutor, (**B**) 100 µM Lonidamine, or (**C**) 10 mM 2-deoxyglucose. Alternatively, cells were equilibrated in 0 mM glucose followed by addition of 2 mM glucose (**D**). Representative examples of the fluorescence change are shown in the inset of the Glutor treatment panel. Closed circles, iG6PSnFR; open circles, iG6PSnFR-D. Each point is the average of N cells, with N listed on the figure.

### Monitoring G6P dynamics in neurons

To evaluate whether iG6PSnFR has an affinity and ΔF/F sufficient to detect physiologically relevant changes in the concentration of G6P in non-immortalized cells, we next tested iG6PSnFR in primary cultured neurons. Just as with immortalized HeLa cells, removal of glucose from the media results in a drop in cytosolic G6P, which can be restored by replenishing glucose (Supplementary Fig. 3A). It has been previously shown that neurons are highly dependent on glycolysis to provide energy during electrical stimulation (*15*). With iG6PSnFR targeted to boutons *via* a fusion with synaptophysin, we observe that a train of 100 stimuli reduced G6P pools (Supplementary Fig. 3B). iG6PSnFR responds as expected, with an immediate drop in fluorescence followed by an approximately 60 second recovery. iG6PSnFR-D has a more complicated fluorescence profile. Fluorescence decreases immediately during the electrical stimulation (potentially an artefact of decreased pH) and then rises, indicating that the concentration of G6P is decreasing with stimulation, as expected. But unlike the G6P-activated sensor, the recovery appears to take place over a much shorter time period. This might be due to the iG6PSnFR-D having higher affinity for G6P than iG6PSnFR and revealing an overshoot of the G6P equilibrium point.

The change in glycolytic metabolite concentrations during and after electrical stimulation was further validated by observing the change in concentration of four molecular species in parallel experiments using iGlucoSnFR2 to observe glucose (*4*), iG6PSnFR to observe glucose-6-phosphate, HYlight to observe fructose-1,6-bisphosphate (FBP) (*16*), and Pegassos to observe pyruvate (*17*) (Fig. 3). Zooming in on the period of stimulation, we see an immediate rise in FBP and concomitant decrease in G6P. There is an immediate but slower decrease in glucose. Once the stimulus is ended, FBP and G6P drift back to their starting equilibrium values, while glucose homeostasis takes longer. Pyruvate increases, but being at the end of the glycolysis, the timing of its accumulation lags behind the consumption of other species.

**Fig. 3.**
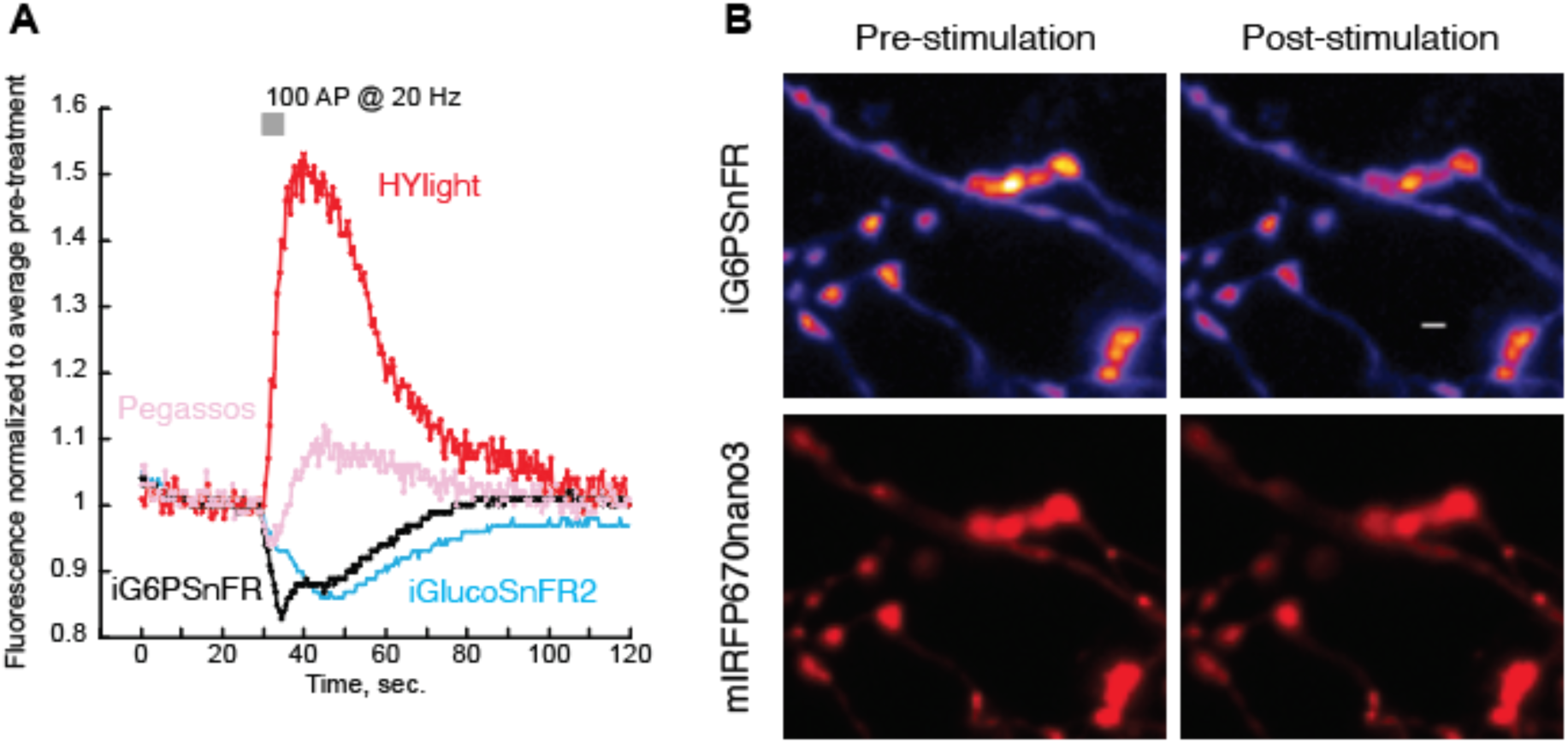
Observation of four different metabolites in parallel experiments in cultured neurons in response to electrical stimulation. (**A**) Sensors for each of the metabolites were anchored to synaptophysin to target them to boutons, facing the cytosolic space. Data points are normalized to the fluorescence intensity just prior to the stimulus. iG6PSnFR.miRFP670nano3 (black, n=12); iGlucoSnFR2.HaloTag-JF635 (blue, n=7); HyLight.HaloTag-JF635 (red, n=8); and Pegassos.HaloTag-JF635 (pink, n=7). (**B**) Representative images of a primary hippocampal neuron expressing synaptophysin targeted iG6PSnFR.miRFP670nano3 construct before and after the stimulus; G6P sensitive channel is pseudo-colored with calibration bar ranging from 0 to 900 AFU; scale bar is 2 µm.

### In vivo imaging of iG6PSnFR in C. elegans neurons

To determine whether neuronal G6P levels change dynamically *in vivo*, we adapted the iG6PSnFR for use in *C. elegans*. For this context we fused iG6PSnFR to mScarlet-I3 (*18*), and expressed it in the ASEL and ASER chemosensory neurons (Fig. 4A), two paired neurons that have been shown to exhibit distinct metabolic profiles (*19*).

**Fig. 4.**
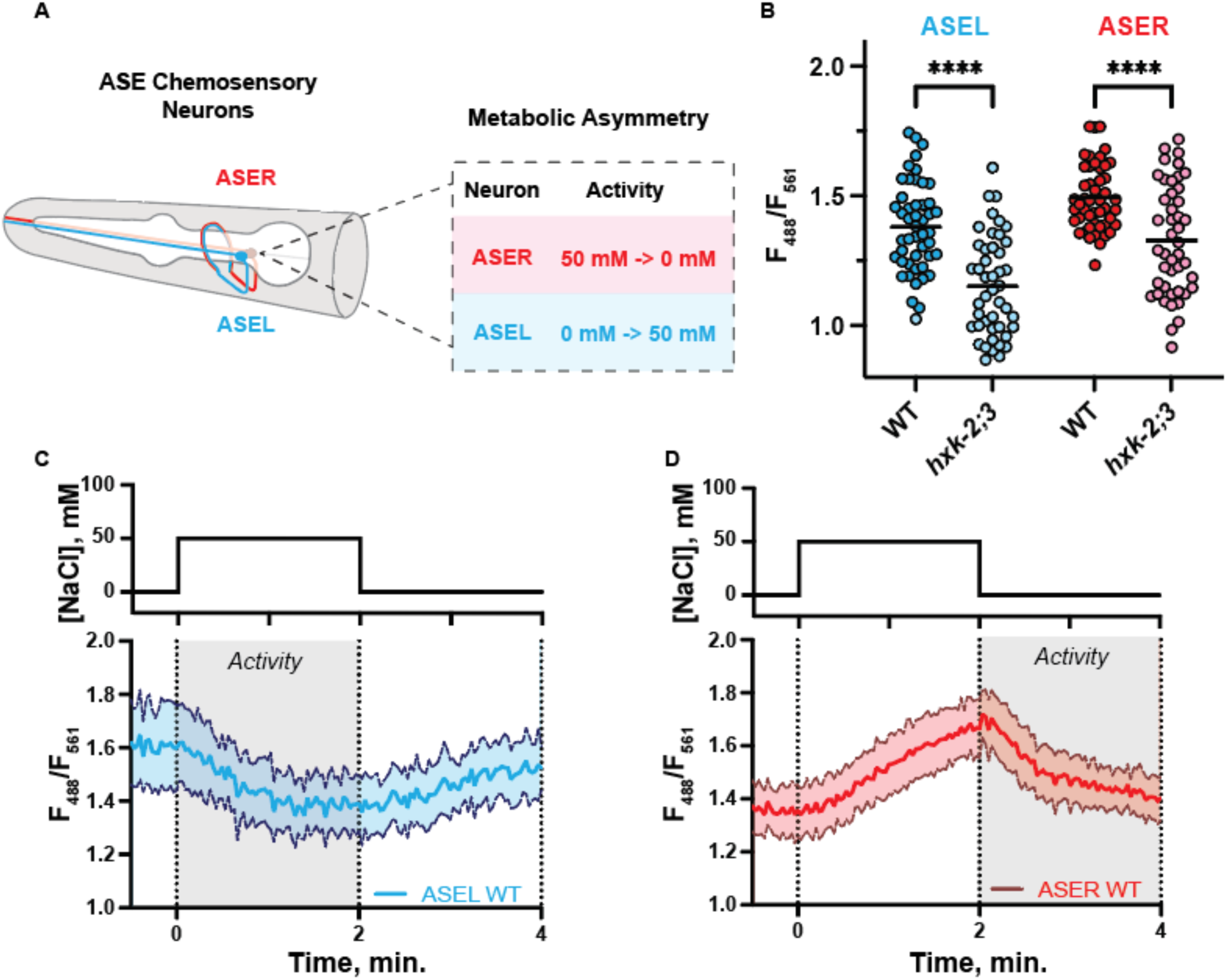
*In vivo* imaging of iG6PSnFR in *C. elegans* ASE sensory neurons. **(A)** Diagram illustrating the bilaterally symmetric, but functionally and metabolically asymmetric, chemosensory neurons ASEL and ASER. (**B**) Mean ratiometric soma iG6PSnFR measurements (relative to mScarlet-I3) in wild type (WT) and *hxk-2;hxk-3* double knockout animals. Significant differences were observed between ASEL WT and ASEL *hxk-2;3* (t(182) = 6.158, p < 0.0001) and ASER WT and ASER *hxk-2;3* (t(182) = 4.471, p < 0.0001). (**C**, **D**) Dynamic iG6PSnFR responses upon a brief pulse of 50 mM NaCl from a 0 mM baseline in either ASEL (**C**) or ASER (**D**) of wild-type animals. Shaded areas around the mean represent the ± 95% confidence interval. Grey shading denotes the windows of neuronal activity for ASEL and ASER, respectively.

We first validated the sensor by observing G6P levels within the cell soma of ASEL and ASER in a hexokinase mutant background. We used CRISPR-Cas9 to generate novel deletion alleles for the genes *hxk-2* and *hxk-3*, the hexokinase orthologs that are principally expressed in these two neurons (*20*). This double mutant displayed significantly reduced G6P levels compared to wild-type animals in both ASEL and ASER (Fig. 4B). Basal G6P levels ratios were also significantly higher in ASER than in ASEL in wild-type animals, consistent with a recent study demonstrating that ASER is more glycolytically active than ASEL (*19*). These results support that iG6PSnFR functions *in vivo* to visualize changing states of G6P.

We next examined whether G6P changed dynamically *in vivo* upon neuronal stimulation, as glycolysis is known from prior literature to be upregulated with neuronal activity (*19*, *21*). ASEL and ASER respond asymmetrically to changes in NaCl concentration: ASEL is stimulated by increases in NaCl levels while ASER is stimulated by decreases in NaCl levels (*22*). We thus observed the responses of iG6PSnFR in these two neurons upon a 50 mM salt pulse stimulation paradigm (Fig 4C, D). When we stimulated ASEL by increasing NaCl from 0 mM to 50 mM, we observed a decrease in the iG6PSnFR/mScarlet-I3 ratio, indicating a decrease in G6P concentration (Fig. 4C). In that same time window, we saw an increase in G6P in ASER while it was inactive (Fig. 4D). When we decreased salt concentration (50 mM to 0 mM) to stimulate ASER, iG6PSnFR signal in ASER sharply decreased whereas the levels in ASEL recovered. These results are similar to the decreases in iG6PSnFR fluorescence observed in cultured neurons upon neuronal stimulation (Fig. 3). Furthermore, these G6P dynamics are consistent with the increases in the downstream metabolite fructose-1,6-bisphosphate in these same neurons (*16*), as those increases occur simultaneously with the use of G6P. These findings strongly demonstrate that iG6PSnFR reports dynamic changes in G6P levels within neurons in both *in culture* and *in vivo* contexts.

### iG6PSnFR reports oscillations of glucose-6-phosphate in mouse islet β-cells

Because of its central position at the entry point of organismal glucose metabolism, monitoring G6P in pancreatic islets can provide valuable insight into how β-cells sense glucose and regulate insulin release under physiological and disease conditions. To that end, we expressed iG6PSnFR-D in mouse islet β-cells islets using an adenovirus containing the insulin promoter. It is well established that β-cell glycolysis initiates a cascade of signaling events that culminate in the release of insulin (*23–25*). In this pathway, a metabolism-driven rise in the ATP/ADP ratio closes ATP-sensitive potassium channels to depolarize the plasma membrane, which opens voltage-dependent calcium channels, triggering Ca^2+^ entry and insulin exocytosis.

To simultaneously monitor G6P dynamics and downstream events in β-cells, we performed three-channel fluorescence imaging. The fluorescence of iG6PSnFR-D was recorded in the green channel, using the shoulder of the GFP excitation/emission spectrum (Ex. 511 nm, Em. 544 nm) to report changes in G6P concentration. The intrinsic fluorescence of NAD(P)H, reflecting the metabolism dependent rise in the redox potential, was simultaneously recorded in the blue channel (Ex. 360 nm, Em. 472 nm). Finally, to monitor the timing of membrane depolarization, we also used Fura Red, a Ca^2+^ sensitive fluorescent dye that has a broad ratiometric excitation profile (Ex. 430 nm and 511 nm), with emission at 650 nm. Observing these three channels in individual islets that were transitioned from 2 mM glucose to 10 mM glucose, we see that G6P and NAD(P)H rise prior to Ca^2+^ influx, as indicated by a decrease in the fluorescence of the iG6PSnFR-D sensor and an increase in NAD(P)H fluorescence (Fig. 5A,B). At steady-state levels of 10 mM glucose, naturally occurring oscillations can be observed (Fig 5C, Supplementary Video S1). The presence of Fura Red does not affect the dynamics of iG6PSnFR-D (Supplementary Fig. S4a), demonstrating compatibility of the biosensor with simultaneous multiplexed imaging.

**Fig. 5.**
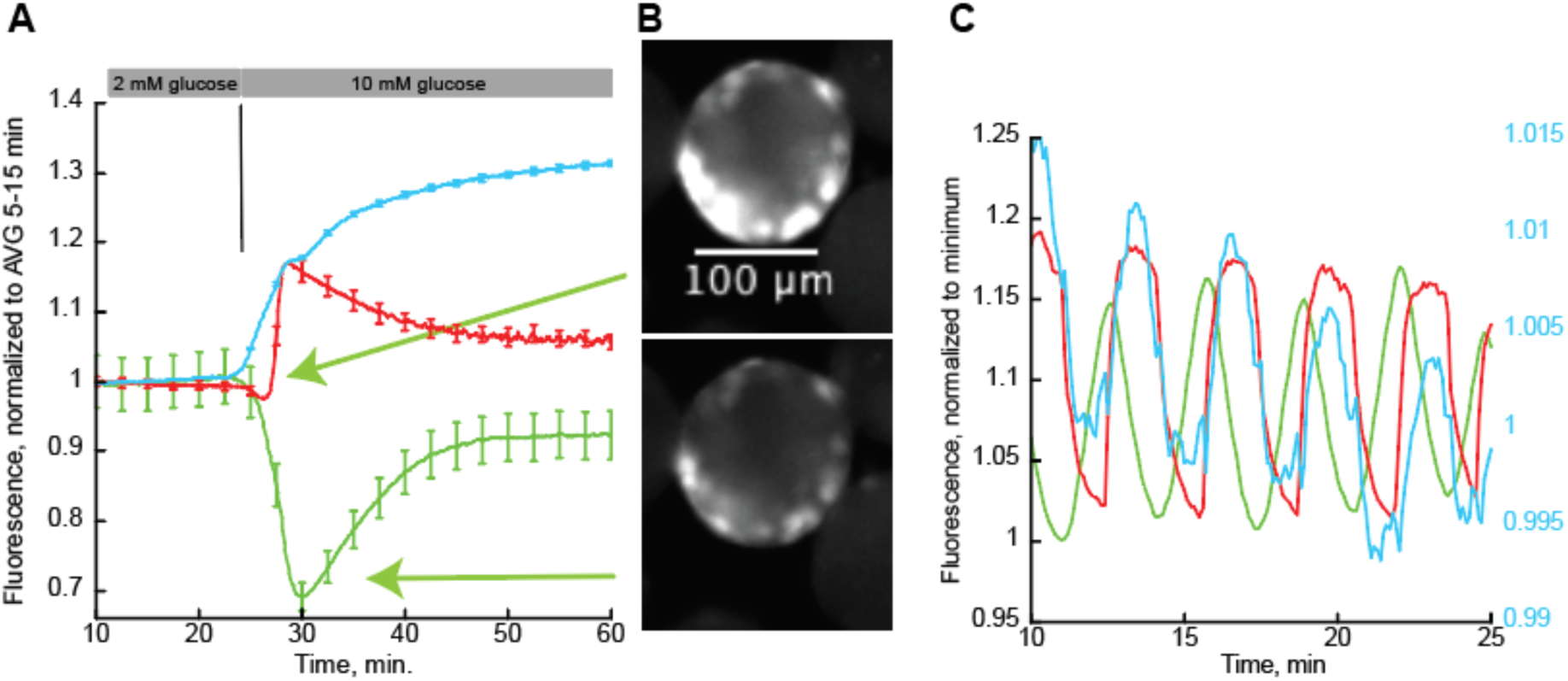
Multiplexed widefield fluorescence imaging of pancreatic islets. (**A**) Fluorescence changes of NAD(P)H via autofluorescence (blue), G6P via iG6PSnFR-D (green), and Ca^2+^ via Fura Red (red). Islets were equilibrated in media containing 2 mM glucose. After 20 minutes, the media was replaced with media containing 10 mM glucose as indicated by the grey horizontal bars and the black vertical line. Fluorescence values (or the ratio of Fura Red fluorescence in the case of Ca^2+^) are normalized to a baseline defined by the average fluorescence during the period of 5 min. to 15 min. Error bars are SEM. N = 50, 50, and 29 islets for the NAD(P)H, G6P, and Ca^2+^, respectively. (**B**) Representative green channel images of an islet expressing iG6PSnFR-D at the brightest and dimmest points of the transition from 2 mM to 10 mM glucose. (**C**) Representative oscillation of the same three species in one islet when the media has remained in 10 mM glucose, normalized to the minimum value.

The G6P-activated variant (iG6PSnFR) was more technically challenging to image than iG6PSnFR-D under widefield microscopy since background fluorescence from flavins overlaps with the GFP excitation/emission spectra (*26*). Since the iG6PSnFR is dim in the unbound state (Supp. Fig. 1), and autofluorescence was high, and we were unable to resolve G6P-dependent fluorescence changes with that variant (Supplementary Fig. S4b). However, when we used 2-channel spinning disk confocal imaging to reduce the contribution of out-of-plane flavin fluorescence (Supplementary Fig. S4c), we were able to observe an increase in G6P induced by elevating media glucose concentration (Supplementary Fig. S4d).

### iG6PSnFR reports differences in steady state G6P concentrations in glycolytic and gluconeogenic hepatocytes intravital imaging

Finally, we used iG6PSnFR to query the steady state distribution of G6P in the livers of live mice (Fig. 6A). The proximity of a hepatocyte to an oxygen-rich portal vein (PV) or an oxygen-depleted central vein (CV) determines whether it is gluconeogenic or glycolytic, respectively (Fig. 6B) (*27*, *28*). However, it is unknown how that balance influences the homeostasis of G6P within those cells. To address that question, and to establish that iG6PSnFR can be used to compare G6P levels across populations of hepatocytes, we expressed both iG6PSnFR variants fused to mIRFP670nano3 in hepatocytes *via* tail-vein injection of AAV harboring the sensor under control of the TBG promoter. The reference reporter miRFP670nano3 was included to normalize for differences in sensor expression arising from the known bias of AAV transduction toward CV hepatocytes in mice (*29*). The differences in intensity of iG6PSnFR relative to mIRFP were clearly visible in the dual-color images (Fig. 6B). When the portal vein (PV) and central vein (CV) were annotated and labeled, the quantification of the green-to-far-red fluorescence intensity ratio (Fig. S5, A and B) showed higher concentrations of G6P in hepatocytes near the CV than near the PV, indicating that glycolytic cells maintain a higher pool of glucose-6-phosphate than gluconeogenic cells. This was true for both iG6PSnFR and iG6PSnFR-D, although the latter expressed less well.

**Fig. 6.**
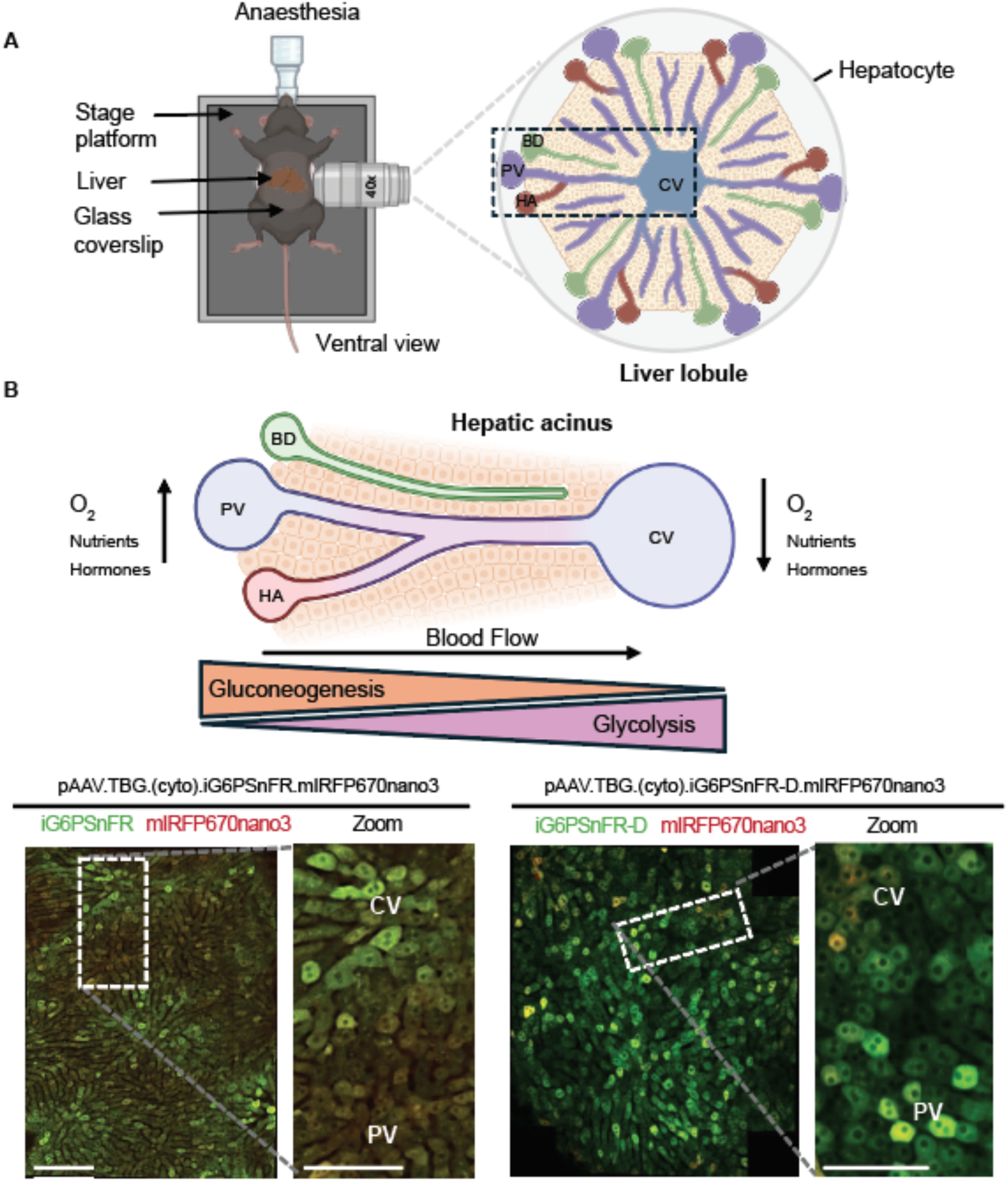
Intravital imaging of iG6PSnFR. (**A**) Experimental setup illustrating intravital imaging of the liver. The zoomed illustration shows a hexagonal liver lobule in which the portal vein (PV), hepatic artery (HA), and bile duct (BD) are located at the corners and the central vein (CV) is located at the center. The liver acinus is highlighted by a dotted box. **(B)** Imaging of iG6PSnFR variants in the liver metabolic gradient. The schematic illustrates unidirectional blood flow in the liver acinus, creating metabolic gradients that dictate the balance between gluconeogenesis and glycolysis along the PV–CV axis. Representative dual-color fluorescence images with iG6PSnFR (sensor) in green and miRFP670nano3 (reference) in red show more G6P (higher green-to-red signal) in CV regions for the G6P-activated sensor. Imaging of iG6PSnFR-D also shows more G6P (lower green-to-red signal) near the CV. Scale bar, 200 μm (100 μm for the insets). Illustrations were created using BioRender.

## Discussion

Here we report the design and validation of a pair of genetically encoded fluorescent sensors for glucose-6-phosphate (G6P). The two sensors’ respond appropriately either positively or negatively in the presence of G6P. The affinity of the sensors appears to be appropriate for observing changes in cytosolic G6P concentrations among multiple preparations and imaging modalities: immortalized cells, primary neurons, soma of neurons in *C. elegans*, isolated pancreatic islets, and intact liver.

G6P lies at a branch point of carbon metabolism where it can proceed through glycolysis, the pentose phosphate pathway, glycogen metabolism and the hexosamine pathway. In combination with other sensors (existent and yet to be created), iG6PSnFR can be used to report not just energetic state, but also which cellular pathways are prioritized. G6P can represent a commitment step where the cell has decided to take up glucose but then must decide whether to push G6P towards creating ATP *via* glycolysis, to invest it to create nucleotides for growth, or NADPH for redox support *via* the pentose phosphate pathway, to store as glycogen or lipids, or to repurpose it for glycosylation via the hexosamine pathway. In cells or tissues that are undergoing glycogenolysis, gluconeogenesis or lipolysis, G6P represents a common intermediate in the utilization of alternative carbon sources in the absence of glucose. In combination with other methods these cellular “decisions” can now be more efficiently monitored. Rerouting metabolism can be directly observed from a variety of cellular perturbations which could include cell signaling, oxidative stress, nutrient availability, drug responses, and disease with temporal resolution on the scale of seconds to minutes.

The application space for iG6PSnFR is large. Recent development of a variety of real time genetically encoded sensors for metabolic intermediates such as glucose (*4*), FBP (*16*, *28*), lactate (*30*), pyruvate (*31*), ATP (*32*), and others are ushering in new opportunities for accessing the role of metabolism in a variety of biologic responses. Traditional metabolomics research relies heavily on LC-MS and radioisotope tracking and can deliver quantitative information regarding a wide range of metabolites (*33*). However, these methods lack spatial and temporal resolution. Regarding spatial resolution, we have demonstrated here that G6P levels vary in regions of the liver adjacent to portal or central veins. Temporal resolution G6P is also provided by neuronal responses to electrical stimulation and relative responses of G6P, FBP, glucose, and pyruvate. In pancreatic islet cells, we were able to resolve changes in G6P, NAD(P)H, and calcium with simultaneous three channel imaging. Such spatial and temporal metabolic heterogeneity is common in other tissues and in cancerous tumors (*34*). These advantages can be extended to identify subcellular distribution differences and dynamics within individual cells. Genetically encoded sensors can be targeted to cytosol, nucleus, synapses, mitochondria, or the ER lumen where there are likely localized metabolic microenvironments that can influence or respond to a variety of cellular stimuli. G6P levels are particularly relevant in the ER lumen. G6P is transported to the ER during glycogenolysis, gluconeogenesis and lipolysis where it is converted to glucose and either used for protein glycosylation or transported back to the cytosol. Defects in ER G6P transport or G6P phosphatase cause glycogen storage disease type 1 (*35*). A direct readout of G6P could also advance therapeutic approaches to this disease.

Metabolic intermediates also serve as allosteric regulators of both metabolic enzyme activity and transcription. For example, G6P regulates the activity of hexokinase (*36*), glycogen synthase (*37*), and glycogen phosphorylase b (*38*). G6P also regulates transcription of genes involved in glucose uptake, glycolysis, and lipogenesis *via* ChREBP and MondaA (*39*, *40*). These two basic helix-loop-helix-leucine zipper transcription factors respond to increased G6P levels, which promotes their translocation to the nucleus and complex formation with MlX. This response to G6P is tissue specific. ChREBP functions mainly in the liver to promote lipid synthesis whereas MondoA predominantly regulates glycolytic genes in skeletal muscle. Temporal and spatial monitoring of G6P represents previously unavailable tool for monitoring these regulatory roles. Creative application of iG6PSnFR should enable a more detailed exploration of this metabolic landscape.

## Materials and Methods

### Mutagenesis & candidate screening & subcloning

The DNA sequence encoding the G6P-binding protein HptA was optimized for expression in mouse using the codon optimization tool of Integrated DNA Technologies (IDT) and cloned into a derivative of the pRSET bacterial expression vector (Invitrogen) *via* BglII and PstI sites. Oligonucleotide directed mutagenesis (*41*) was used to introduce NheI and AgeI sites (encoding AS and TG) at the positions chosen for insert of cpSFGFP. cpSFGFP was amplified by PCR from iGlucoSnFR2 with primers including NheI and AgeI sites at the 5’ and 3’ ends and inserted into the HptA gene by restriction digest. Primers encoding libraries of the residues at the junction of the binding protein and the cpSFGFP were used to make libraries of variants using the uracil template method (*41*) and transformed into T7 Express cells (New England Biolabs). Individual colonies were picked into single wells of a 96-well plate (Arctic White Square Well Polypropylene 2.2mL AWLS-T110-10) containing 850 µL autoinduction media (*42*) and 100 µg/mL ampicillin and set to shake for 18 hours at 30°C at 350 rpm. Bacterial pellets were collected by centrifuging the 96-well plates at 3000 RPM in a swinging bucket rotor. Supernatant was removed, replaced with 500 µL Tris-buffered saline (TBS), and vortexed to resuspend the bacterial pellets. The collection and resuspension process was performed three times to remove endogenous G6P. Pellets were frozen at-20°C overnight. Clarified lysate was obtained by thawing the pellets with in 1 mL PBS, resuspending by vortex, and subsequent centrifugation. 100 µL of lysate was transferred to black flat bottom 96-well plates by multi-channel pipet. GFP fluorescence was measured in a Tecan Safire 2 fluorescence plate reader (excitation 485/20 nm, emission 535/20 nm), G6P added to 2 mM, and then measured again. Candidates with higher than starting point ΔF/F were isolated, sequenced, and carried forward for additional mutation and screening.

### Protein purification and in vitro characterization

Bacterial expression plasmids encoding iG6PSnFR proteins were transformed into T7 express cells. Individual colonies were used to inoculate 300 mL autoinduction media with 100 µg/mL ampicillin in a 2L flask and grown at 30°C for 18 hours at 225 RPM. Cells were pelleted by centrifugation at 2000g and resuspended in 30 mL TBS and frozen at - 20°C overnight. Bacterial suspension was thawed in room temperature water, sonicated for 5 minutes (5 sec on, 5 sec off) on ice. Cellular debris was removed by centrifugation at 6000g for 10 min. Supernatant was transferred to a clean tube and clarified by centrifugation at 35000g for 60 min. Clarified lysate was loaded onto a 5 mL IMAC Fast Flow column (Cytiva) at 2 mL/min, rinsed with 40 mL PBS, and eluted with a gradient to 150 mM imidazole over 60 min. Fluorescent green fractions were pooled, concentrated by centrifugal ultrafiltration, and dialyzed with TBS to remove any residual G6P or inorganic phosphate. Protein concentration was determined by diluting 2 µL protein into 18 µL 1 M NaOH, measuring absorbance at 447 nm, and using the extinction coefficient of GFP = 44,000 µM^-1^ cm^-1^.

All *in vitro* characterization was performed with 0.2 µM protein in TBS at room temperature. G6P titrations were performed in a Tecan Safire 2 plate reader (excitation 485/20 nm, emission 535/20 nm), by making serial dilutions of a concentration stock of 1 M glucose and adding them to 100 µL purified protein. Excitation spectra were collected by observing emission at 515/5 nm while varying excitation wavelength (5 nm bandpass) from 300 to 500 nm. Emission spectra were collected by exciting at 485/5 nm and observing emission (5 nm bandpass) from 500 to 600 nm. Kinetic measurements were made by mixing equal volumes protein with glucose in a SX-20 stopped flow fluorimeter (Applied Photophysics) 5 times and averaging the traces.

### Immortalized cell culture

Cell culture maintenance and electroporation were performed by the Howard Hughes Medical Institute at the Janelia Research Campus, Immortalized Cell Line Culture Shared Resource Core Facility (RRID:SCR_026515). HeLa cells (ATCC, CCL-2) were cultured in DMEM supplemented with 10% FBS, 1% L-glutamine, and 1% penicillin/streptomycin, and incubated in a CO₂ incubator at 37°C with 5% CO₂. For transfection, the reaction mixture was prepared by combining 1 × 10⁶ HeLa cells with 0.5 µg of each plasmid per reaction in 20 µL of electroporation SE solution (Lonza, V4SC-1096). Each reaction was transferred to a 96-well electroporation plate, and electroporation was performed according to the manufacturer’s protocol. Subsequently, 2.5 × 10⁵ cells per well were plated in a 24-well plate and further incubated until imaging.

After cells were transfected and given 18 hours to recover and express recombinant sensor proteins, HaloTag ligand-coupled JFX650 was added to a final concentration of 100 nM in Mammalian Cell Imaging buffer (20 mM HEPES pH 7.5, 120 mM NaCl, 2.5 mM KCl, 2 mM CaCl_2_, 2 mM MgCl_2_) with 2 mM glucose. We imaged cells in a Cytation multi-well plate reader using both the GFP channel (to detect G6P or glucose) and the CY5 channel to detect HaloTag coupled to the far-red fluorophore JFX650 to normalize for variable protein expression. Where indicated, an equal volume of buffer was added with 2x the final concentration of drug.

### Primary neuronal culture

All animal-related experiments were performed in accordance with protocols approved by the Weill Cornell Medicine IACUC, protocols number 0601-450A and 2009-0026. Wild-type rats were of the Sprague-Dawley strain (Charles River Laboratories, strain code: 400, RRID: RGD_734476). Hippocampal CA1-CA3 neurons were isolated from 1- to 2-day-old rats of mixed gender, plated on poly-ornithine-coated coverslips, transfected 7 days after plating, and imaged 14-21 days after plating as previously described (*43*).

### Live-cell imaging of primary neurons

Live-cell imaging was performed as previously described (*44*). All imaging experiments were performed on a custom-built laser illuminated epifluorescence Zeiss Axiovert 200 microscope with Andor iXon camera (model: DU-897U-CS0-BVF) and 40X 1.3 NA Fluar Zeiss objective. Cells were maintained at 37 ^0^C and constantly perfused at a rate of 0.1 ml/min with a Tyrode’s solution containing 5 mM glucose, 119 mM NaCl, 2.5 mM KCl, 2 mM CaCl_2_, 2 mM MgCl_2_, 50 mM HEPES (pH 7.4). The buffer was supplemented with 0.01 mM 6-cyano-7-nitroquinoxalibe-2, 3-dione (CNQX) and 0.05 mM D,L-2-amino-5-phosphonovaleric acid (APV) to suppress post-synaptic responses. Action potentials were evoked by passing 1-ms current pulses, yielding fields of approximately 10 V cm^-1^ *via* platinum-iridium electrodes. A385 High Current Stimulus Isolator from World precision Instruments was used. For all imaging experiments, background-subtracted images were used for further analysis.

### C. elegans - Sensor adaptation and cloning

The iG6PSnFR sensor was first codon-optimized and synthesized for *C. elegans* expression using the GeneArt Strings DNA Synthesis Service from ThermoFisher. HaloTag was replaced with mScarlet-I3 (*18*) as the reference fluorophore, enabling ratiometric imaging by comparing emission with excitation by 488 nm or 561 nm light. To examine G6P dynamics in a cell-type specific manner, we expressed the adapted sensor using the *flp-6* promoter for cell-specific expression in both ASEL and ASER neurons. ASER-specific expression of TagBFP was used to differentiate the two neurons (*gcy-5p*::TagBFP). Plasmid construction was carried out using Gibson Assembly, and transgenic animals were generated by gonadal microinjection (*45*) using standard techniques.

### C. elegans - CRISPR-Cas9 genome editing

We generated *hxk-2(ola589)* and *hxk-3(ola590)* single mutants by CRISPR-mediated full gene deletion, following established injection protocols (*46*). Guide RNAs flanking the full gene sequence were identified using the CRISPOR web tool (*47*) and the IDT online gRNA design platform. Guides were synthesized as Edit-R Modified Synthetic crRNAs by Horizon Discovery (Dharmacon). The *dpy-10(cn64)* allele was co-targeted to allow identification of potentially edited F1 animals by the roller phenotype. Single-stranded homology repair templates with 35 base pair overlaps flanking each cut site were synthesized by the Keck Oligonucleotide Synthesis Facility at Yale School of Medicine. All guide RNA and homology template sequences are listed in Supplementary Table S1. *hxk-2(ola589)* deletions were generated using IGCR16 and IGCR17 as flanking guides with IGHT13 as the repair template. *hxk-3(ola590)* deletions were generated using IGCR18 and IGCR19 with IGHT14 as the repair template. Single mutants were crossed together to generate the *hxk-2; hxk-3* double mutant, which was subsequently crossed into the *flp-6p::iG6PSnFR* sensor strain.

### C. elegans - Baseline imaging

Basal iG6PSnFR measurements were performed using a hypoxia microfluidics device as previously described (*48*), with compressed air flowing throughout the duration of the experiment to maintain normoxic conditions. Animals were immobilized in standard imaging buffer (50 mM NaCl, 5 mM levamisole, 25 mM potassium phosphate pH 6, 1 mM calcium chloride, 1 mM magnesium sulfate) for 10 minutes prior to imaging to allow animals to relax onto their dorsal/ventral sides and visualize both ASE neurons simultaneously.

### C. elegans - Salt stimulation

Salt stimulation experiments were adapted from Ref. (*19*). To measure glycolytic responses to sensory-evoked neuronal activity animals were loaded into a microfluidics device under active flow of 0 mM NaCl imaging buffer and allowed to equilibrate for 3 minutes prior to imaging, under active flow of 0 mM NaCl buffer. Animals were then subjected to a 5-minute stimulus protocol consisting of a 1-minute baseline at 0 mM NaCl, a 2-minute pulse of 50 mM NaCl, and a 2-minute return to 0 mM NaCl. Under this paradigm, ASEL is excited by the onset of the NaCl stimulus while ASER is excited by its removal, enabling simultaneous measurement of ON and OFF sensory responses within the same animal (*22*).

### C. elegans - Microscopy and Analysis

Imaging was performed on a Nikon Ti2 + CSU-W1 spinning disk confocal microscope with a Hamamatsu Orca-Fusion BT CMOS camera at 16-bit pixel depth using a 4x objective. iG6PSnFR was excited at 488 nm (30% laser power, 100 msec exposure) and mScarlet-I3 was excited at 561 nm (5% laser power, 100 msec exposure). Salt stimulation measurements were acquired at 0.5 Hz at a resolution of 1024 x 1024 pixels. Mean soma fluorescence values were quantified for each neuron across the recording and reported as raw ratiometric values (F₄₈₈/F₅₆₁). Statistical analysis and data visualization were performed using GraphPad Prism. Differences in basal iG6PSnFR ratios across groups were assessed using an ordinary one-way ANOVA with Holm-Šídák’s multiple comparisons test. Image analyses for both experiments were performed in Fiji (*49*).

### Imaging of β-cell targeted iG6PSnFR in mouse pancreatic islets

Full-length adenoviruses containing the G6P sensors iG6PSnFR and iG6PSnFR-D were commercially prepared by cloning into an adenovirus serotype 5 vector (VectorBuilder) downstream of the rat insulin promoter and β-globin intron to facilitate β-cell specific expression in isolated mouse islets, as previously described (*24*). For widefield imaging of G6P and cytosolic Ca^2+^, islets were preincubated in 2.5 µM Fura Red (Molecular Probes F3020) in RPMI 1640 for 45 min at 37°C prior to being placed in a glass-bottomed imaging chamber (Warner Instruments) on Nikon Ti2 microscope equipped with a 10×/0.5NA SuperFluor objective (Nikon). Islets expressing the G6P biosensors were alternately barcoded by preincubation with 2 µmol/L DiR (Thermo Fisher Scientific D12731) in 1 mL islet media for 10 min at 37°C to facilitate simultaneous imaging of both biosensors. The chamber was perfused with standard imaging solution (in mM: 135 NaCl, 4.8 KCl, 2.5 CaCl_2_, 1.2 MgCl_2,_ 20 HEPES, 0.5 glutamine) with glucose indicated in the figure legends. Temperature was maintained at 33°C with an SF-28 inline solution heater and QE-1 chamber heater (Warner Instruments), and the flow rate was set to 0.25 mL/min (Fluigent). Excitation was provided by a SOLA SEII 365 (Lumencor) attenuated with an ND8 neutral density filter (Nikon). Fluorescence emission was collected with a Hamamatsu ORCA-Flash4.0 V2 Digital CMOS camera every 6 s. A single region of interest was used to quantify the average response of each islet using NIS-Elements software (Nikon Instruments). Excitation (Ex) or emission (Em) filters (ET type; Chroma Technology) were used in combination with an FF459/526/596-Di01 dichroic beamsplitter (Semrock) with wavelengths listed in nm/bandpass as follows: iG6PSnFR, Ex 511/20, Em 544/24; Fura Red, Ex 434/20 and Ex 511/20x, Em 650/60 (Ratio 434/511); and NAD(P)H, Ex 360/20, Em 472/30. Spinning disk confocal imaging was conducted on Nikon CSU-W1 Spinning Disk Confocal Microscope equipped with a 10×/0.5NA SuperFluor objective (Nikon Instruments). iG6PSnFR and Fura Red were excited at 488 nm (1-2.5% power), emission was split with a 561LP dichroic, filtered (Em 531/50, iG6PSnFR; Em 691/64, Fura Red), and collected with Hamamatsu Orca Quest2 cameras using 4×4 binning. A second camera was used to image Fura Red fluorescence at 488 nm excitation and 691 nm emission.

### Intravital imaging of liver hepatocytes

Six-weeks-old male C57BL/6J mice (strain #000664) purchased from Jackson were housed under a 12-hour light/dark cycle with ad libitum access to food and water. At ten weeks of age, mice were injected via the lateral tail vein with approximately 4 × 10¹¹ adeno-associated virus serotype 8 (AAV8) particles. AAVs encoding either the G6P-activated sensor iG6PSnFR (AAV.TBG.(cyto).iG6PSnFR.mIRFP670nano3) or the G6P-deactivated sensor iG6PSnFR-D (AAV.TBG.(cyto).iG6PSnFR-D.mIRFP670nano3) were administered to separate animals. Intravital imaging of the liver was performed 3 days after AAV administration. Imaging was conducted between 9:00 am ± 2 hours for both sensors. All animal procedures were carried out in accordance with NIH guidelines and were approved by the Institutional Animal Care and Use Committee (Protocol #25-0280) at Janelia Research Campus, Howard Hughes Medical Institute.

Images were collected using a Leica Stellaris 8 confocal microscope equipped with a 40 x oil-immersion objective, using a pinhole size of 2 Airy units, a zoom factor of 2, and an image resolution of 1024 × 1024 pixels. Sensor (iG6PSnFR) and reference (miRFP670nano3) fluorescence images were acquired in the green and far-red channels and used to calculate the sensor-to-reference fluorescence intensity ratio in Fiji. Portal vein (PV) and central vein (CV) regions were manually annotated during image acquisition based on blood flow visualization and morphological features observed under the microscope. Large regions of liver containing multiple lobules were acquired in each animal.

## Supporting information

SuppFigs

## Acknowledgments

JSM and TAB are members for the Tool Translation Team at Janelia Research Campus.

## Author contributions

Conceptualization: JSM

Methodology: JSM, ADW, IJG, ZK, RA

Investigation: JSM, ZK, AT, IJG, SMH, ZK, RA

Visualization: AT, IJG, SMH, ZK, RA

Supervision: TAB, TAR, DAC-R, MJM, DF

Writing—original draft: JSM, TAB

Writing—review & editing: All authours

## Data and materials availability

All data are available in the main text or the supplementary materials.

All DNA constructs (including permutations of sequence elements not used in this publication) are being made available on Addgene (Plasmid ####).

## References

1. F. Rajas, A. Gautier-Stein, G. Mithieux, Glucose-6 Phosphate, a Central Hub for Liver Carbohydrate Metabolism. Metabolites 9, 282 (2019).

2. W. W. Chen, E. Freinkman, T. Wang, K. Birsoy, D. M. Sabatini, Absolute Quantification of Matrix Metabolites Reveals the Dynamics of Mitochondrial Metabolism. Cell 166, 1324–1337.e11 (2016).

3. I. Pérez-Chávez, E. H. Gilglioni, D. Ezeriņa, J. Messens, E. N. Gurzov, Single-cell imaging of liver metabolic dynamics using fluorescent biosensors. Trends Endocrinol. Metab., doi: 10.1016/j.tem.2025.10.003 (2025).

4. J. S. Marvin, P. Mächler, C. Meng, T. Ates, R. H. Patel, R. Adhikari, M. A. Makurath, Z. Ku, D. Feliciano, D. Atasoy, G. Cui, D. Kleinfeld, T. A. Brown, iGlucoSnFR2: A genetically encoded fluorescent sensor for measuring intracellular or extracellular glucose in vivo in mouse brain. Sci. Adv. 11, eadz3889 (2025).

5. M. Wang, Q. Guo, K. Zhu, B. Fang, Y. Yang, M. Teng, X. Li, Y. Tao, Interface switch mediates signal transmission in a two-component system. Proc. Natl. Acad. Sci. 117, 30433–30440 (2020).

6. B. Sit, S. M. Crowley, K. Bhullar, C. C.-L. Lai, C. Tang, Y. Hooda, C. Calmettes, H. Khambati, C. Ma, J. H. Brumell, A. B. Schryvers, B. A. Vallance, T. F. Moraes, Active Transport of Phosphorylated Carbohydrates Promotes Intestinal Colonization and Transmission of a Bacterial Pathogen. PLoS Pathog. 11, e1005107 (2015).

7. J. S. Marvin, Y. Shimoda, V. Magloire, M. Leite, T. Kawashima, T. P. Jensen, I. Kolb, E. L. Knott, O. Novak, K. Podgorski, N. J. Leidenheimer, D. A. Rusakov, M. B. Ahrens, D. M. Kullmann, L. L. Looger, A genetically encoded fluorescent sensor for in vivo imaging of GABA. Nat. Meth. 16, 763–770 (2019).

8. K. Davidsen, J. S. Marvin, A. Aggarwal, T. A. Brown, L. B. Sullivan, An engineered biosensor enables dynamic aspartate measurements in living cells. eLife 12, RP90024 (2024).

9. J. S. Marvin, B. Scholl, D. E. Wilson, K. Podgorski, A. Kazemipour, J. A. Müller, S. Schoch, F. J. U. Quiroz, N. Rebola, H. Bao, J. P. Little, A. N. Tkachuk, E. Cai, A. W. Hantman, S. S.-H. Wang, V. J. DePiero, B. G. Borghuis, E. R. Chapman, D. Dietrich, D. A. DiGregorio, D. Fitzpatrick, L. L. Looger, Stability, affinity, and chromatic variants of the glutamate sensor iGluSnFR. Nat. Methods 15, 936–939 (2018).

10. P. C. Rosen, S. Sullere, P. Fu, J. R. Martínez-François, D. J. Brooks, E. Kim, C. Gu, G. Yellen, Inorganic phosphate and the rapid mobilization of metabolic energy in neurons. Proc. Natl. Acad. Sci. 123 (2026).

11. J. B. Grimm, L. Xie, J. C. Casler, R. Patel, A. N. Tkachuk, N. Falco, H. Choi, J. Lippincott-Schwartz, T. A. Brown, B. S. Glick, Z. Liu, L. D. Lavis, A General Method to Improve Fluorophores Using Deuterated Auxochromes. JACS Au 1, 690–696 (2021).

12. E. S. Reckzeh, G. Karageorgis, M. Schwalfenberg, J. Ceballos, J. Nowacki, M. C. M. Stroet, A. Binici, L. Knauer, S. Brand, A. Choidas, C. Strohmann, S. Ziegler, H. Waldmann, Inhibition of Glucose Transporters and Glutaminase Synergistically Impairs Tumor Cell Growth. Cell Chem. Biol. 26, 1214–1228.e25 (2019).

13. K. Nath, L. Guo, B. Nancolas, D. S. Nelson, A. A. Shestov, S.-C. Lee, J. Roman, R. Zhou, D. B. Leeper, A. P. Halestrap, I. A. Blair, J. D. Glickson, Mechanism of antineoplastic activity of lonidamine. Biochim. Biophys. Acta (BBA) - Rev. Cancer 1866, 151–162 (2016).

14. W. Chen, M. Guéron, The inhibition of bovine heart hexokinase by 2-deoxy-d-glucose-6-phosphate: characterization by 31P NMR and metabolic implications. Biochimie 74, 867–873 (1992).

15. A. C. Kokotos, A. M. Antoniazzi, S. R. Unda, M. S. Ko, D. Park, D. Eliezer, M. G. Kaplitt, P. D. Camilli, T. A. Ryan, Phosphoglycerate kinase is a central leverage point in Parkinson’s disease–driven neuronal metabolic deficits. Sci. Adv. 10, eadn6016 (2024).

16. J. N. Koberstein, M. L. Stewart, C. B. Smith, A. I. Tarasov, F. M. Ashcroft, P. J. S. Stork, R. H. Goodman, Monitoring glycolytic dynamics in single cells using a fluorescent biosensor for fructose 1,6-bisphosphate. Proc National Acad Sci 119, e2204407119 (2022).

17. K. Harada, T. Chihara, Y. Hayasaka, M. Mita, M. Takizawa, K. Ishida, M. Arai, S. Tsuno, M. Matsumoto, T. Ishihara, H. Ueda, T. Kitaguchi, T. Tsuboi, Green fluorescent protein-based lactate and pyruvate indicators suitable for biochemical assays and live cell imaging. Sci. Rep. 10, 19562 (2020).

18. T. W. J. Gadella, L. van Weeren, J. Stouthamer, M. A. Hink, A. H. G. Wolters, B. N. G. Giepmans, S. Aumonier, J. Dupuy, A. Royant, mScarlet3: a brilliant and fast-maturing red fluorescent protein. Nat. Methods 20, 541–545 (2023).

19. A. D. Wolfe, L. Niu, L. S. Yilmaz, S. Ravikumar, M. Thomas, A. J. M. Walhout, Z.-W. Wang, R. H. Goodman, D. A. Colón-Ramos, Glycolytic Specialization Shapes Neuronal Physiology and Function in vivo. bioRxiv, 2026.02.17.706437 (2026).

20. S. R. Taylor, G. Santpere, A. Weinreb, A. Barrett, M. B. Reilly, C. Xu, E. Varol, P. Oikonomou, L. Glenwinkel, R. McWhirter, A. Poff, M. Basavaraju, I. Rafi, E. Yemini, S. J. Cook, A. Abrams, B. Vidal, C. Cros, S. Tavazoie, N. Sestan, M. Hammarlund, O. Hobert, D. M. Miller, Molecular topography of an entire nervous system. Cell 184, 4329–4347.e23 (2021).

21. C. M. Díaz-García, R. Mongeon, C. Lahmann, D. Koveal, H. Zucker, G. Yellen, Neuronal Stimulation Triggers Neuronal Glycolysis and Not Lactate Uptake. Cell Metab. 26, 361–374.e4 (2017).

22. H. Suzuki, T. R. Thiele, S. Faumont, M. Ezcurra, S. R. Lockery, W. R. Schafer, Functional asymmetry in Caenorhabditis elegans taste neurons and its computational role in chemotaxis. Nature 454, 114–117 (2008).

23. M. J. Merrins, B. E. Corkey, R. G. Kibbey, M. Prentki, Metabolic cycles and signals for insulin secretion. Cell Metab. 34, 947–968 (2022).

24. S. L. Lewandowski, R. L. Cardone, H. R. Foster, T. Ho, E. Potapenko, C. Poudel, H. R. VanDeusen, S. M. Sdao, T. C. Alves, X. Zhao, M. E. Capozzi, A. H. de Souza, I. Jahan, C. J. Thomas, C. S. Nunemaker, D. B. Davis, J. E. Campbell, R. G. Kibbey, M. J. Merrins, Pyruvate Kinase Controls Signal Strength in the Insulin Secretory Pathway. Cell Metab. 32, 736–750.e5 (2020).

25. T. Ho, E. Potapenko, D. B. Davis, M. J. Merrins, A plasma membrane-associated glycolytic metabolon is functionally coupled to KATP channels in pancreatic α and β cells from humans and mice. Cell Rep. 42, 112394 (2023).

26. A. C. Croce, G. Bottiroli, Autofluorescence spectroscopy and imaging: a tool for biomedical research and diagnosis. Eur. J. Histochem. 58, 2461 (2014).

27. T. M. Yasaka, C. K. Kim, V. Meadows, S. P. Monga, Zonation, Zonation, Zonation: The Real Estate of the Liver. Annu. Rev. Pathol.: Mech. Dis. 21, 185–212 (2026).

28. J. Tyler, A. A. Vishwanath, T. Menon, T. Duarah, R. Adhikari, J. N. Koberstein, D. Feliciano, I. Espinosa-Medina, D. Colon-Ramos, A. G. Tebo, Improved sensors for fructose-1,6-bisphosphate enable in vivo imaging of glycolysis. bioRxiv, 2026.04.29.721630 (2026).

29. P. Bell, L. Wang, G. Gao, M. E. Haskins, A. F. Tarantal, R. J. McCarter, Y. Zhu, H. Yu, J. M. Wilson, Inverse zonation of hepatocyte transduction with AAV vectors between mice and non-human primates. Mol. Genet. Metab. 104, 395–403 (2011).

30. S. Hario, G. N. T. Le, H. Sugimoto, K. Takahashi-Yamashiro, S. Nishinami, H. Toda, S. Li, J. S. Marvin, S. Kuroda, M. Drobizhev, T. Terai, Y. Nasu, R. E. Campbell, High-Performance Genetically Encoded Green Fluorescent Biosensors for Intracellular l-Lactate. ACS Cent. Sci. 10, 402–416 (2024).

31. R. Arce-Molina, F. Cortés-Molina, P. Y. Sandoval, A. Galaz, K. Alegría, S. Schirmeier, L. F. Barros, A. S. Martín, A highly responsive pyruvate sensor reveals pathway-regulatory role of the mitochondrial pyruvate carrier MPC. Elife 9, e53917 (2020).

32. J. S. Marvin, A. C. Kokotos, M. Kumar, C. Pulido, A. N. Tkachuk, J. S. Yao, T. A. Brown, T. A. Ryan, iATPSnFR2: A high-dynamic-range fluorescent sensor for monitoring intracellular ATP. Proc. Natl. Acad. Sci. 121, e2314604121 (2024).

33. C. Jang, L. Chen, J. D. Rabinowitz, Metabolomics and Isotope Tracing. Cell 173, 822–837 (2018).

34. J. Kim, R. J. DeBerardinis, Mechanisms and Implications of Metabolic Heterogeneity in Cancer. Cell Metab. 30, 434–446 (2019).

35. J. Y. Chou, H. S. Jun, B. C. Mansfield, Type I glycogen storage diseases: disorders of the glucose-6-phosphatase/glucose-6-phosphate transporter complexes. J. Inherit. Metab. Dis. 38, 511–519 (2015).

36. T. Ureta, P. A. Lazo, A. Sols, Allosteric inhibition of brain hexokinase by glucose 6-phosphate in the reverse reaction. Arch. Biochem. Biophys. 239, 315–319 (1985).

37. M. Bouskila, R. W. Hunter, A. F. M. Ibrahim, L. Delattre, M. Peggie, J. A. van Diepen, P. J. Voshol, J. Jensen, K. Sakamoto, Allosteric Regulation of Glycogen Synthase Controls Glycogen Synthesis in Muscle. Cell Metab. 12, 456–466 (2010).

38. S. Aiston, B. Andersen, L. Agius, Glucose 6-Phosphate Regulates Hepatic Glycogenolysis Through Inactivation of Phosphorylase. Diabetes 52, 1333–1339 (2003).

39. C. W. Peterson, C. A. Stoltzman, M. P. Sighinolfi, K.-S. Han, D. E. Ayer, Glucose Controls Nuclear Accumulation, Promoter Binding, and Transcriptional Activity of the MondoA-Mlx Heterodimer. Mol. Cell. Biol. 30, 2887–2895 (2010).

40. C. A. Stoltzman, C. W. Peterson, K. T. Breen, D. M. Muoio, A. N. Billin, D. E. Ayer, Glucose sensing by MondoA:Mlx complexes: A role for hexokinases and direct regulation of thioredoxin-interacting protein expression. Proc. Natl. Acad. Sci. 105, 6912–6917 (2008).

41. T. A. Kunkel, Rapid and efficient site-specific mutagenesis without phenotypic selection. Proc. Natl. Acad. Sci. 82, 488–492 (1985).

42. F. W. Studier, Protein production by auto-induction in high-density shaking cultures. Protein Expr. Purif. 41, 207–234 (2005).

43. R. J. Farrell, A. C. Kokotos, T. A. Ryan, Primary hippocampal and cortical neuronal culture and transfection. doi: 10.17504/protocols.io.ewov1qxr2gr2/v1 (2023).

44. C. Pulido, T. A. Ryan, Protocol for Neuronal Live-imaging of primary cultures. doi: 10.17504/protocols.io.q26g7pn4qgwz/v1 (2023).

45. L. R. Girard, T. J. Fiedler, T. W. Harris, F. Carvalho, I. Antoshechkin, M. Han, P. W. Sternberg, L. D. Stein, M. Chalfie, WormBook: the online review of Caenorhabditis elegans biology. Nucleic Acids Res. 35, D472–D475 (2007).

46. A. Paix, A. Folkmann, G. Seydoux, Precision genome editing using CRISPR-Cas9 and linear repair templates in C. elegans. Methods 121, 86–93 (2017).

47. J.-P. Concordet, M. Haeussler, CRISPOR: intuitive guide selection for CRISPR/Cas9 genome editing experiments and screens. Nucleic Acids Res. 46, W242–W245 (2018).

48. S. Jang, Z. Xuan, R. C. Lagoy, L. M. Jawerth, I. J. Gonzalez, M. Singh, S. Prashad, H. S. Kim, A. Patel, D. R. Albrecht, A. A. Hyman, D. A. Colón-Ramos, Phosphofructokinase relocalizes into subcellular compartments with liquid-like properties in vivo. Biophys. J. 120, 1170–1186 (2021).

49. J. Schindelin, I. Arganda-Carreras, E. Frise, V. Kaynig, M. Longair, T. Pietzsch, S. Preibisch, C. Rueden, S. Saalfeld, B. Schmid, J.-Y. Tinevez, D. J. White, V. Hartenstein, K. Eliceiri, P. Tomancak, A. Cardona, Fiji: an open-source platform for biological-image analysis. Nat. Methods 9, 676–682 (2012).

