## Supplementary material for "iG6PSnFR: A genetically encoded fluorescent sensor for observing glucose-6-phosphate dynamics in living preparations": SuppFigs

Jonathan S. Marvin*, *et al.*

**This PDF file includes:**

Figs. S1 to S5
Tables S1 to S2

**Other Supplementary Materials for this manuscript include the following:**
Movies S1

### Supplementary Figures

#### Fig. S1a.

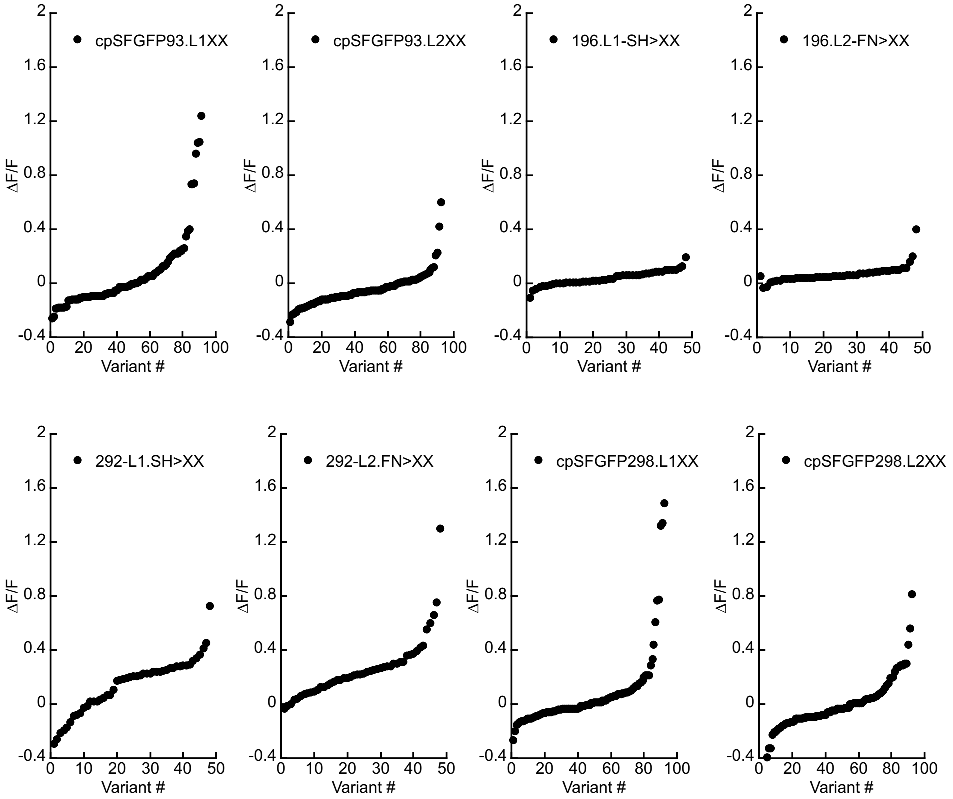


**Supp. Fig S1a.** *Sensor design and in vitro characterization.* (**a**) Initial screening of potential for fluorescence changes with four different insertion sites of cpSGFP into HptA. Residues of the fluorescent protein at the junction of the binding protein were randomized (L1.SH>XX or L2.FN>XX) and 48 or 96 variants were screened by measuring GFP fluorescence of clarified bacterial lysate before and after addition of G6P to 2 mM in TBS.

**Fig. S1b.**

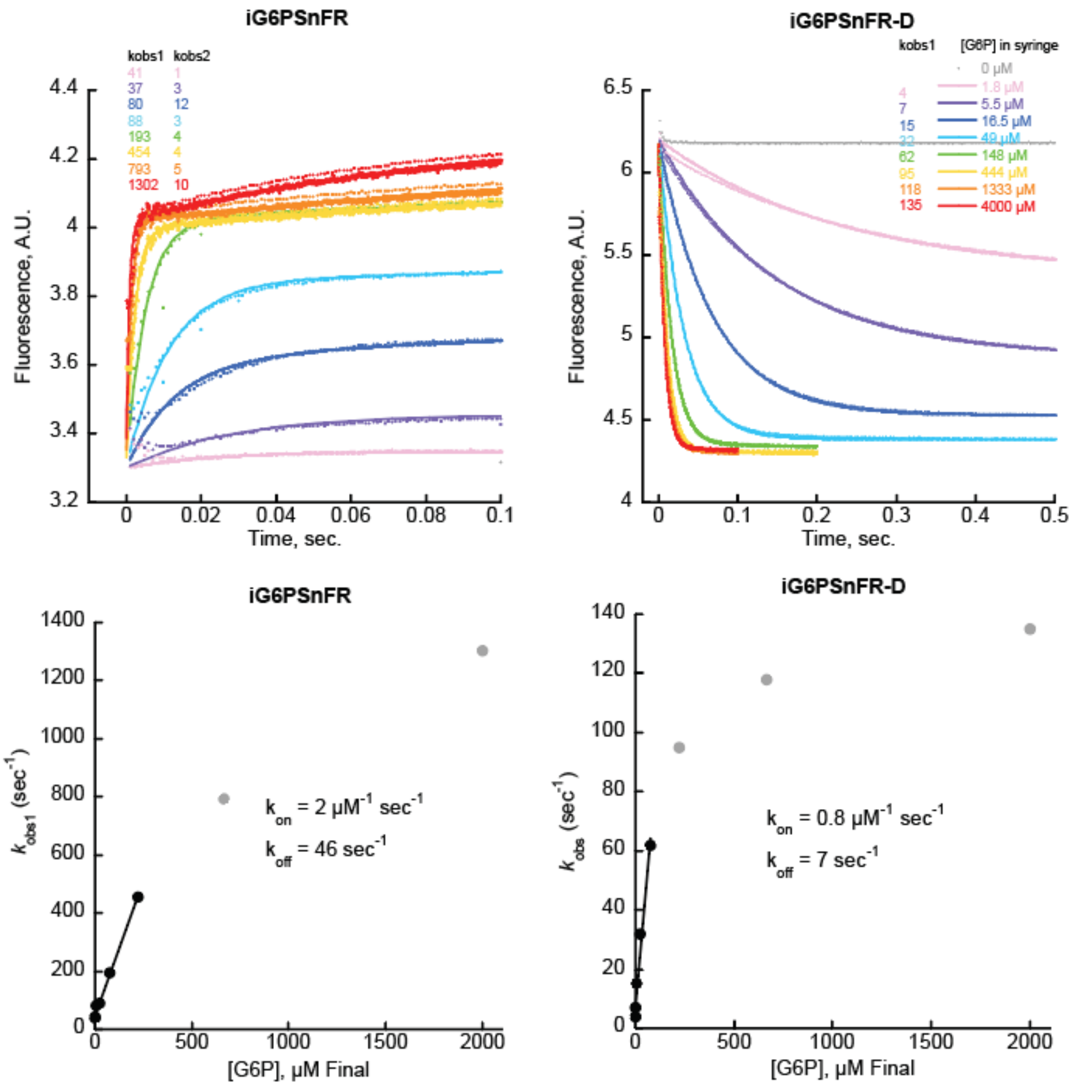


**Supp. Fig S1b.** *Sensor design and in vitro characterization*. (**b**) Binding kinetics of sensors iG6PSnFR and iG6PSnFR-D as determined by stopped-flow fluorescence. Purified sensor proteins were dialyzed into TBS to remove any endogenous G6P. Equal volumes of 0.2 µM sensor and G6P (concentrations listed in figure legend) were mixed in an Applied Photophysics SX20 stopped flow fluorimeter with 490 nm excitation LED and 525 nm longpass emission filter. Top: Rate of change fit well to a single exponential for iG6PSnFR-D, but iG6PSnFR only fits well once a second exponential rise component was added to the curve fit equation. Bottom: Observed rates ($k_{obs}$) were plotted versus final concentration of G6P (half that of the concentration of G6P in the syringe). Fitting the linear portion (black data points) yields $k_{on}$ and $k_{off}$.

**Fig. S1c.**

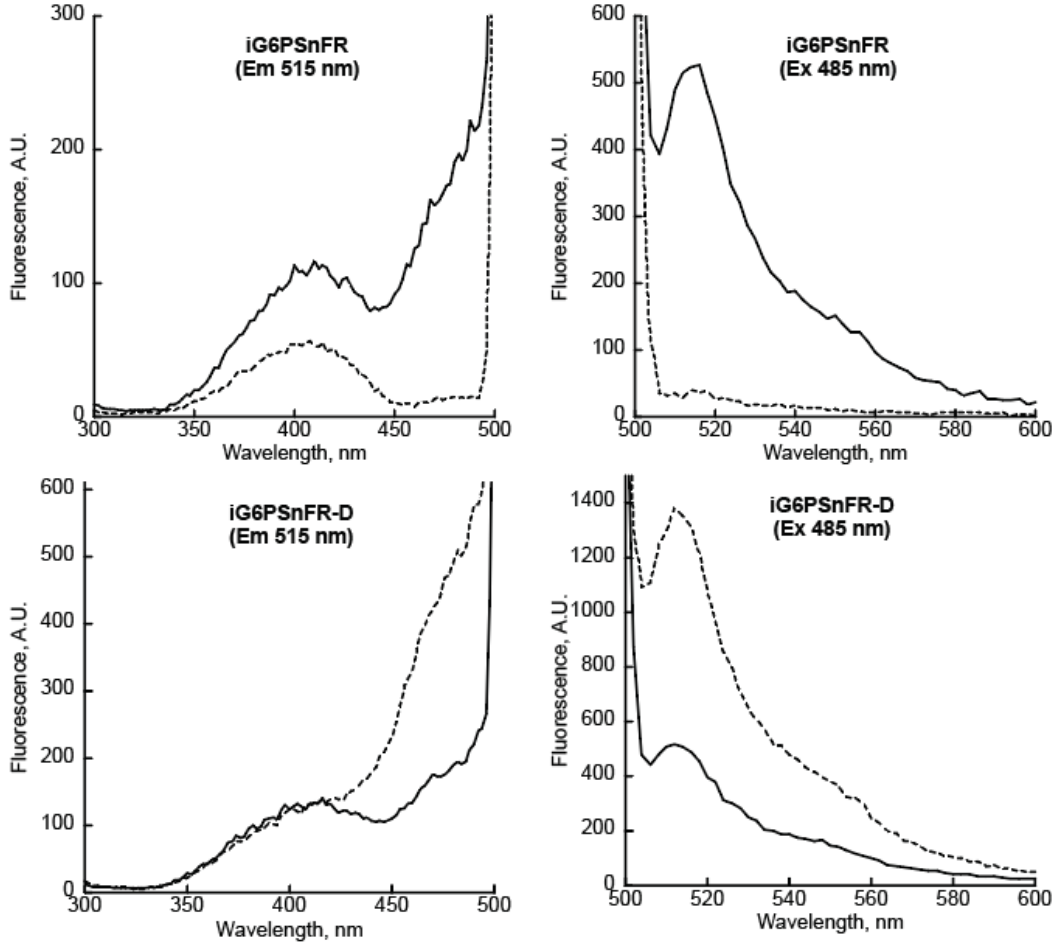


**Supp. Fig S1c.** *Sensor design and in vitro characterization*. (**c**) Spectral properties of the sensors using typical GFP parameters. Spectra of 0.2 µM iG6PSnFR protein (top) or iG6PSnFR-D (bottom) in TBS without G6P (dashed lines) or with 10 mM G6P (solid lines). All spectra were collected with 5 nm bandpass. Excitation spectra were collected observing emission at 515 nm. Emission spectra were collected with excitation at 485 nm.

**Fig. S1d.**

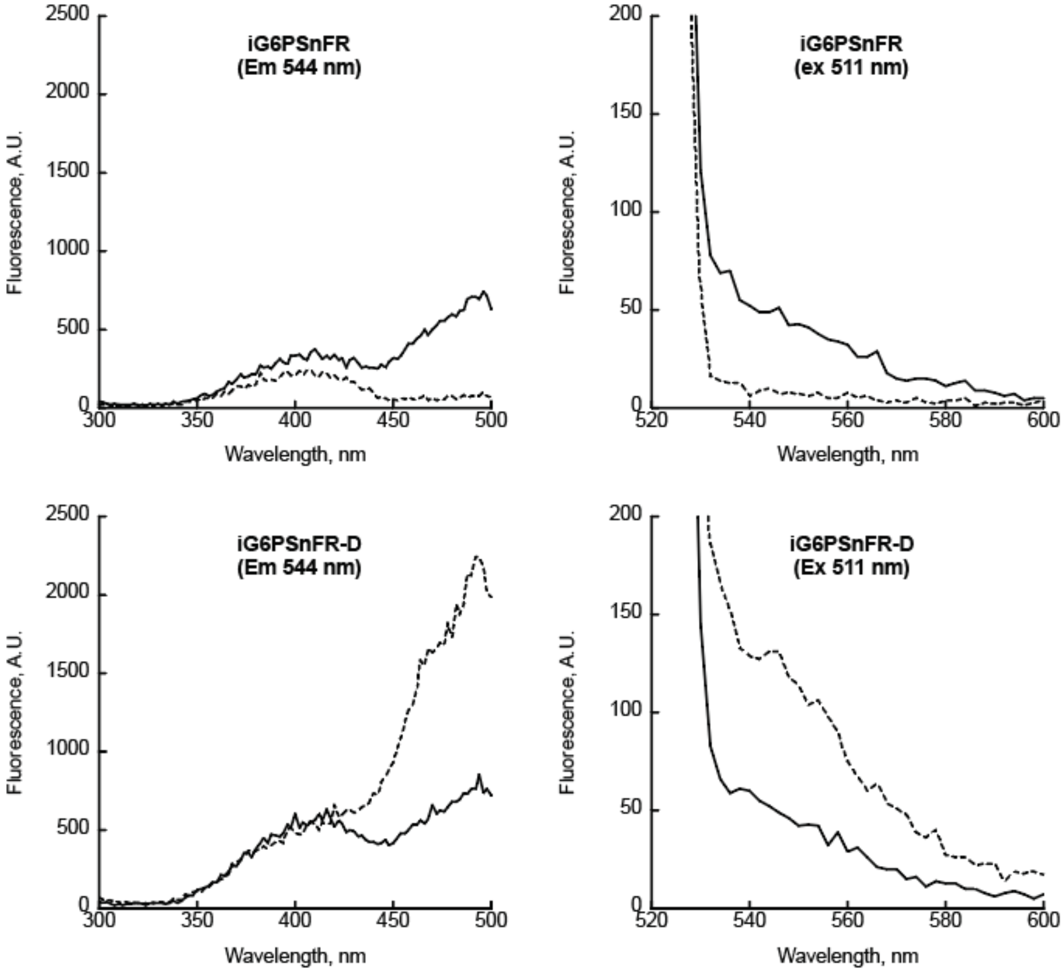


**Supp. Fig S1d.** *Sensor design and in vitro characterization*. (**d**) Spectral properties of the sensors using typical YFP parameters. Spectra of 0.2 μM iG6PSnFR protein (top) or iG6PSnFR-D (bottom) in TBS without G6P (dashed lines) or with 10 mM G6P (solid lines). All spectra were collected with 5 nm bandpass. Excitation spectra were collected observing emission at 544 nm. Emission spectra were collected with excitation at 511 nm.

**Fig. S1e.**

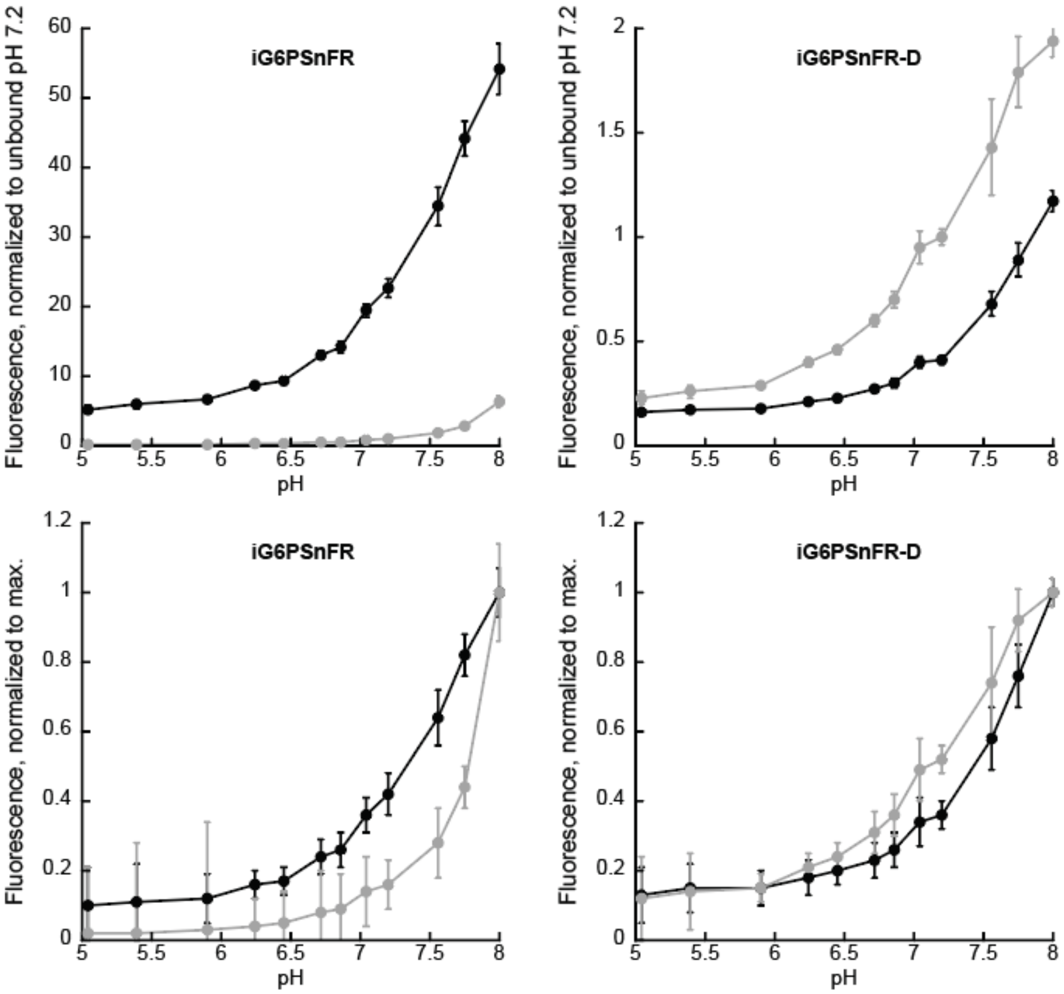


**Supp. Fig S1e.** *Sensor design and in vitro characterization*. (**e**) pH profiles of sensors. Each sensor was diluted to 0.2 µM in 1 mL TBS buffer with the pH adjusted to between 5 and 8 (12 different pH values, x-axis). Those solutions were aliquoted into 6 wells (100 µL each) of a 96-well plate (making a 6 x12 grid of samples for each protein). Fluorescence was measured, G6P added to 2 mM (10 uL from a stock of 20 mM G6P made up in the corresponding pH adjusted buffer), and measured again. Top: Fluorescence values were normalized to the average fluorescence of solutions without G6P at pH 7.2. Bottom: To better illustrate the differences in pH dependence of the bound and unbound states, fluorescence values were normalized to the maximum fluorescence intensity (pH 8.00). All measurements were performed in TBS with 485 nm excitation (7.5 nm bandpass) and 515 nm emission (15 nm bandpass). Data points are average of six technical replicates ± s.d. (define black and gray traces here)

**Fig. S1f.**

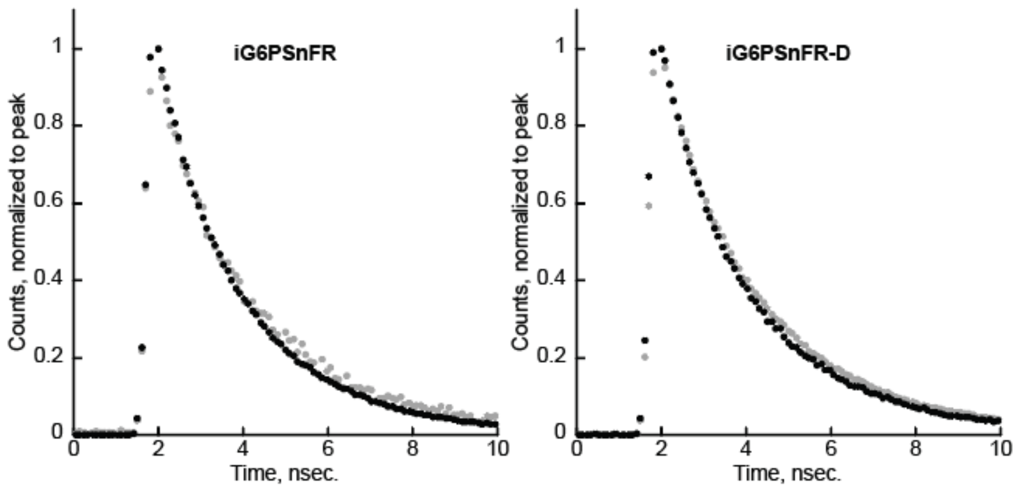


**Supp. Fig S1f.** *Sensor design and in vitro characterization*. (**f**) Fluorescence lifetime properties. The fluorescence lifetime of each sensor (1 µM in TBS) was measured on a Leica SP8 confocal microscope with 488 nm excitation and 515-545 nm emission. Data points are average of three technical replicates in the absence of G6P (grey) or with 10 mM G6P (black). Single exponent decays yield the following lifetimes (nsec.): iG6PSnFR unbound 2.32 ± 0.03, bound 2.12 ± 0.01. iG6PSnFR-D unbound 2.44 ± 0.01, bound 2.28 ± 0.01.

**Fig. S1g.**

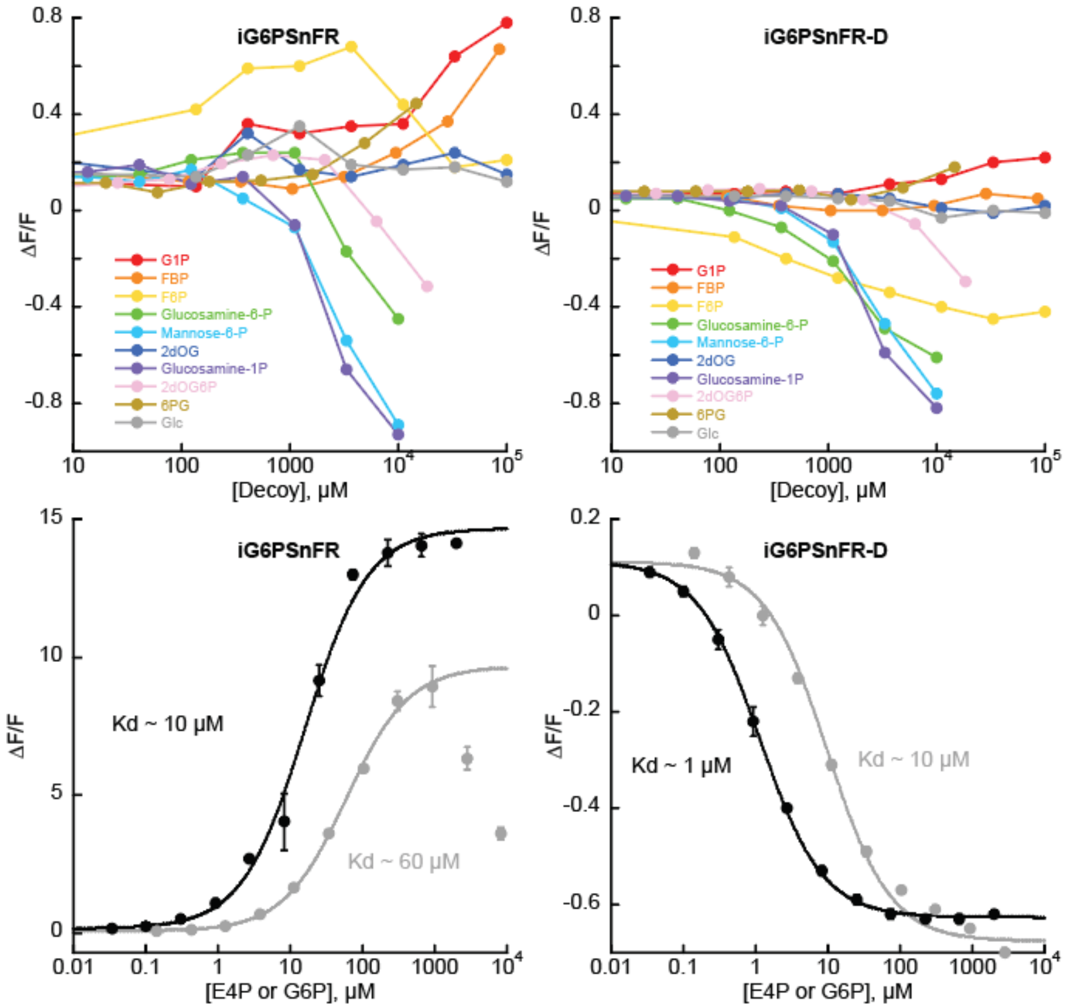


**Supp. Fig S1g.** *Sensor design and in vitro characterization*. **(g)** Ligand specificity. 0.2 μM sensor protein in TBS was titrated with a variety of decoys. Top: glucose-1-phosphate, fructose-1,6-bisphosphate, fructose-6-phosphate, and 6-phosphogluconate, 2-deoxyglucose (an inhibitor of hexokinase) and its metabolic product 2-deoxyglucose-6-phosphate, and glucose show no appreciable change in fluorescence up to concentrations as high as 10 mM. Phosphorylated decoys glucosamine-6-phosphate, mannose-6-phosphate, and glucosamine-1-phosphate showed a decrease in fluorescence at concentrations above 1 mM. These compounds were only available in small quantities and dissolution in TBS yielded highly acidic solutions. The volumes of those stocks were too small to neutralize pH, even with a pH meter with a small probe. Bottom: both sensors appear to have measurable affinity for the phosphorylated four-carbon carbohydrate erythrose-4-phosphate (grey). All measurements were performed in TBS with 485 nm excitation (7.5 nm bandpass) and 515 nm emission (15 nm bandpass). Data points are average of three technical replicates ± s.d..

**Fig. S1h.**

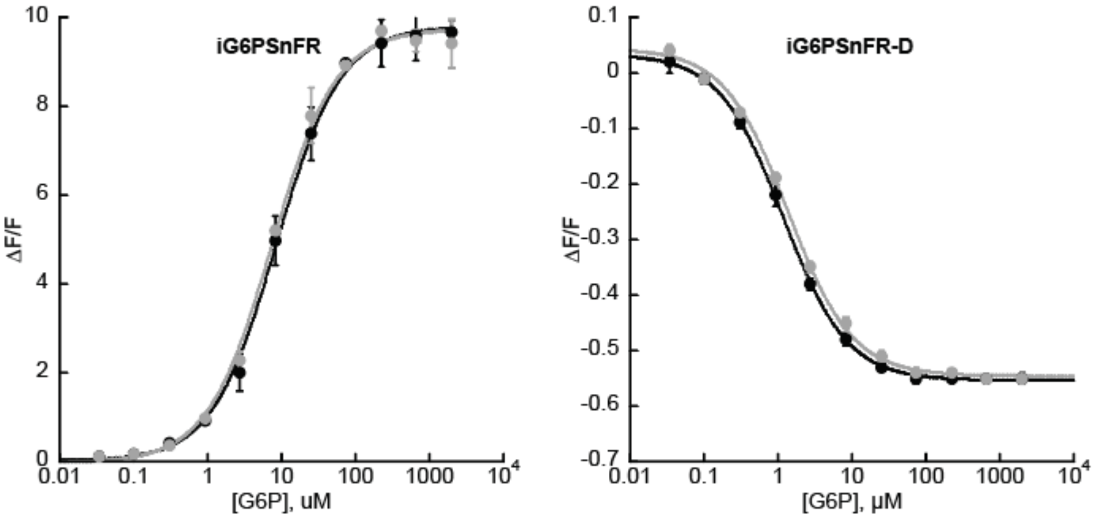


**Supp. Fig S1h.** *Sensor design and in vitro characterization*. (**h**) The affinity of iG6PSnFR for G6P is not affected by glucose. 0.2 µM sensor protein in TBS was titrated with G6BP in the absence of glucose (black) or in the presence of 10 mM glucose (grey). All measurements were performed in TBS with 485 nm excitation (7.5 nm bandpass) and 515 nm emission (15 nm bandpass). Data points are average of three technical replicates ± s.d..

**Fig. S1i.**

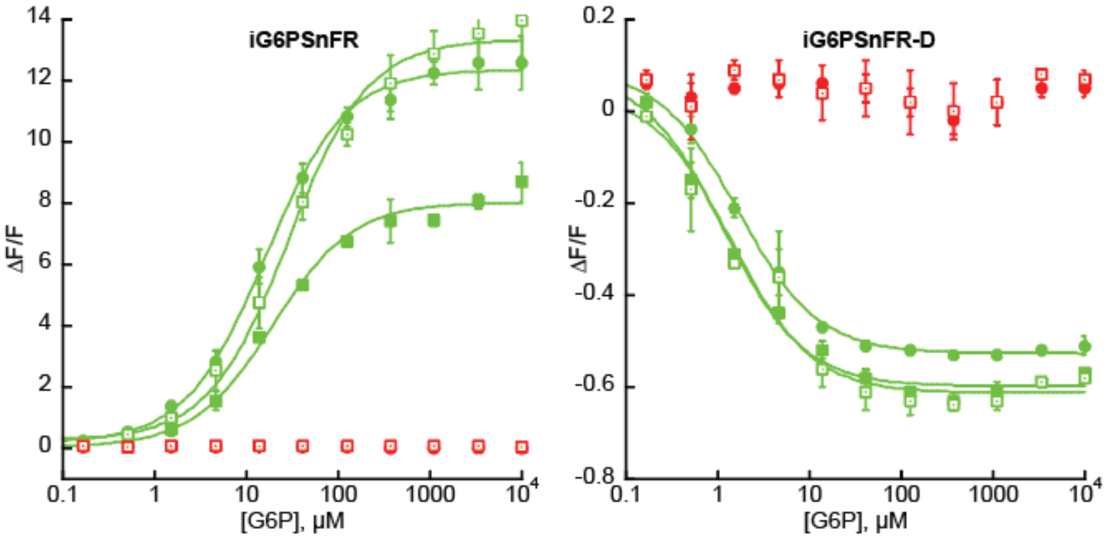


**Supp. Fig S1i.** *Sensor design and in vitro characterization*. (**i**) The affinity of iG6PSnFR for G6P is not affected by C-terminal fusions of HaloTag (solid circles) or mScarlet-I3 (open squares) or mRuby3 (closed squares). All measurements were performed in TBS with 485 nm excitation (7.5 nm bandpass) and 515 nm emission (15 nm bandpass) for the green channel and 560 nm excitation (7.5 nm bandpass) and 600 nm emission (15 nm bandpass) for the red channel. Data points are average of four technical replicates ± s.d..

**Fig. S2.**

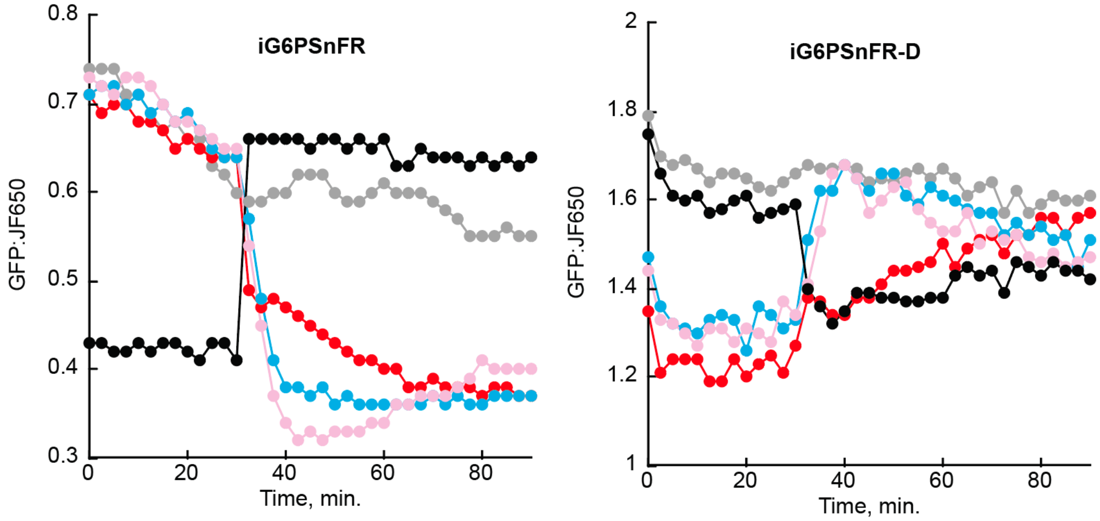


**Supp. Fig S2**. **Effect of metabolic perturbations on G6P levels in HeLa cells.** HeLa cells transfected with (cyto).iG6PSnFR.HaloTag or (cyto).iG6PSnFR-D.HaloTag and labelled with JFX650 were equilibrated with HEPES buffer containing 2 mM glucose and imaged in a Cytation multi-well plate imager. After 30 minutes (as indicated by the arrows), drugs were added to final concentrations of 10 µM Glutor (blue), 100 µM Lonidamine (pink), or 10 mM 2-deoxyglucose (red). Alternatively, cells were equilibrated in 0 mM glucose followed by addition of 2 mM glucose (black). Left, iG6PSnFR; right, iG6PSnFR-DThis is the same data as in Fig. 2 but grouped by sensor instead of treatment.

**Fig. S3.**

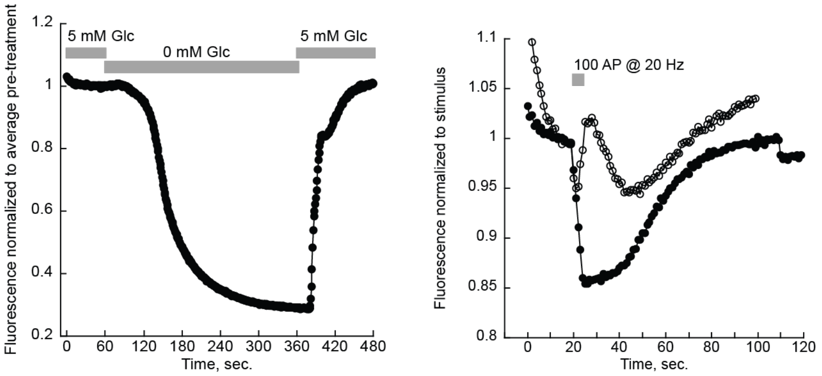


**Supp Fig S3**. *Validation in neuronal culture*. (a) Cultured primary neurons expressing iG6PSnFR.miRFP670nano3. At the times indicated about the graph, the media was exchanged for media containing 0 mM glucose and green fluorescence dropped after a short lag. Fluorescence from the iG6PSnFR sensor is restored when 5 mM glucose is added back to the media. Data points are average of four neurons. (b) Cultured primary neurons expressing either iG6PSnFR (solid circles) or the iG6PSnFR-D (open circles) and stimulated for 5 seconds at the point indicated. Data points are average of 13 and 3 neurons, respectively.

**Fig. S4.**

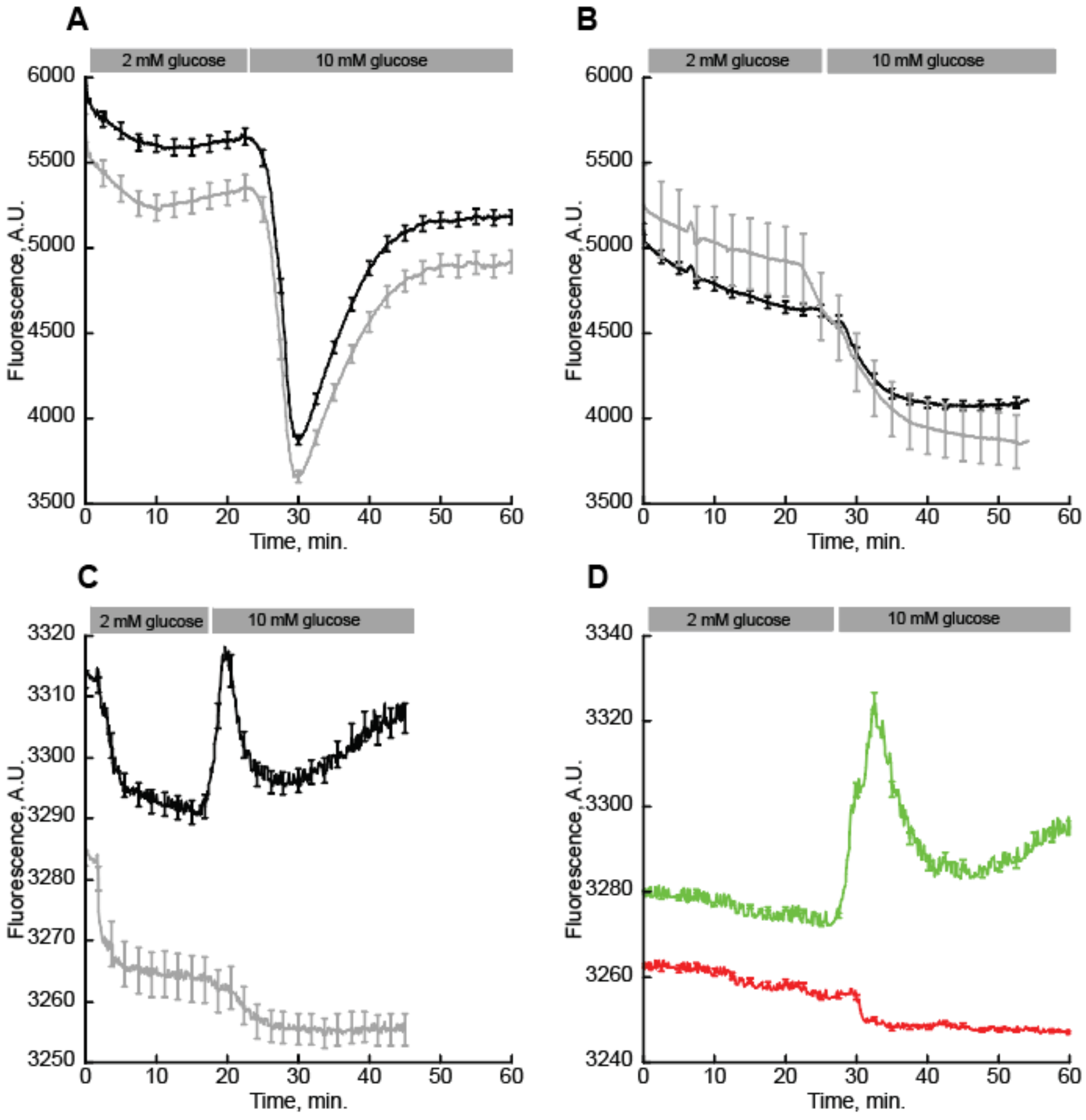


**Supp. Fig. S4.** *Imaging of pancreatic islets in response to glucose elevation from 2 mM to 10 mM*. (a) Widefield imaging of iG6PSnFR-D fluorescence (Ex 475 nm, Em 544 nm) with Fura Red (black, N = 29) or without Fura Red (grey, N = 21). (b) Widefield imaging of iG6PSnFR fluorescence (black, N = 17) or infected islets (grey, N = 4). (c) Spinning disk confocal imaging (Ex 488 nm, Em 531 nm) of iG6PSnFR (black, N = 11) or uninfected (grey, N = 6) islets. (d) Spinning disk confocal two-channel imaging of iG6PSnFR (green, Ex 488 nm, Em 531 nm, N = 9) with FuraRed (red, Ex 488 nm, Em 691). Fura Red is imaged only in the free form. Spinning disk confocal images collected with 2 μm step size, 40 μm range. Error bars are SEM.

**Fig. S5.**

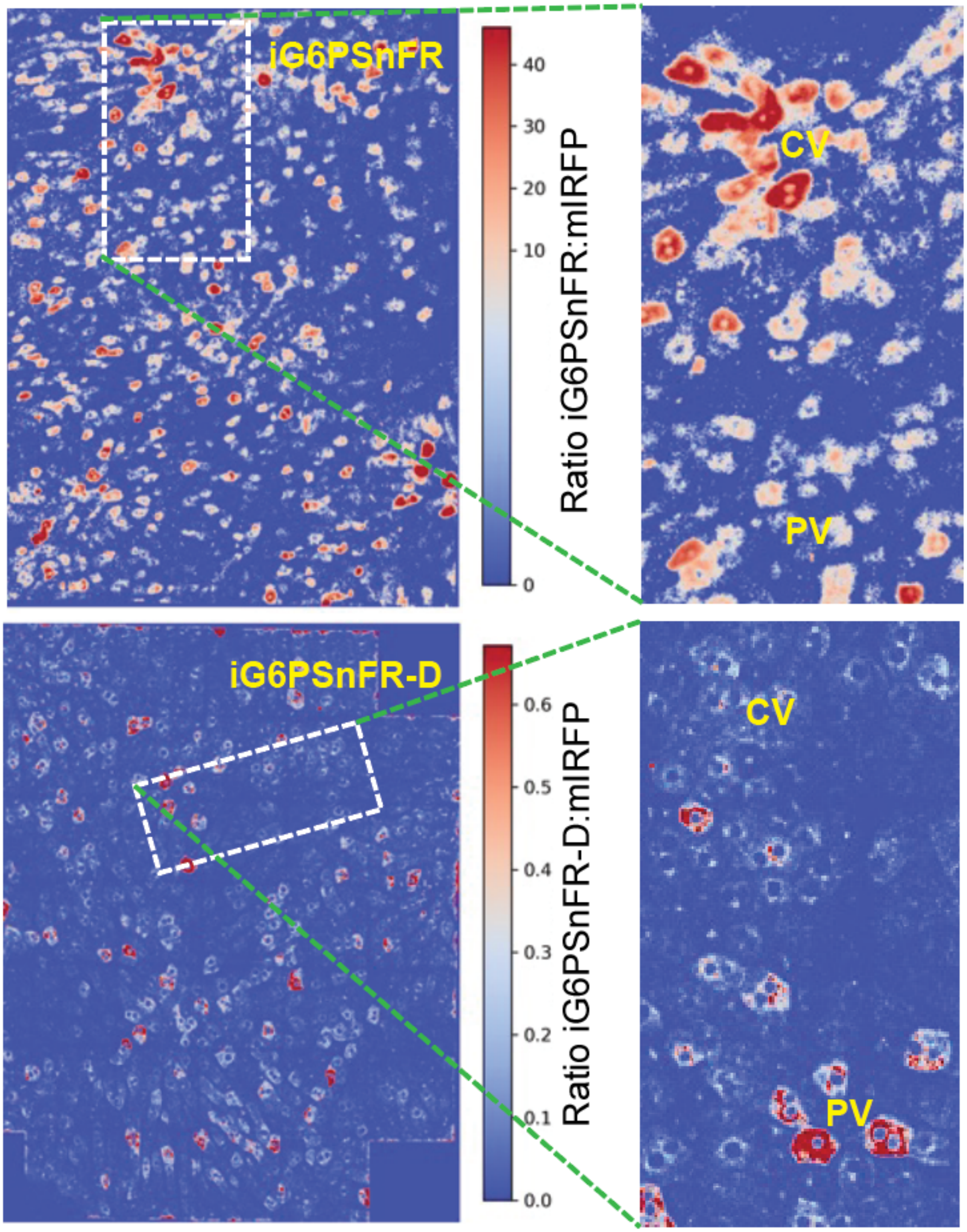


**Supp. Fig. S5**. *Intra-vital imaging of the liver.* Heat maps showing the ratio of iG6PSnFR fluorescence to the mIRFP reference channel (to control for expression artefacts.)

**Table S1.** *C. elegans* CRISPR guide RNAs and repair templates.

| Name | Sequence | Target description/Allele description |
|---|---|---|
| IGCR16 | gcgtaattgggcttacaaag | gRNA targeting the 3' end of *hxk-2* for complete gene deletion |
| IGCR17 | TCGAGCTCGGCATTCAATAT | gRNA targeting the 5' end of *hxk-2* for complete gene deletion |
| IGHT13 | ctacacaaaATGCTGGGTATCATCGAGCTCGGCATTCAgtaagcccaattacgcccccacaaattgccatatacg | *hxk-2* homology repair template; full gene deletion repair template |
| IGCR18 | ACGTGCCTATACTGATGATG | gRNA targeting the 3' end of *hxk-3* for complete gene deletion |
| IGCR19 | GGTTCTACCACCAGAAGTCG | gRNA targeting the 5' end of *hxk-3* for complete gene deletion |
| IGHT14 | cagtaatctccacaaaaATGAAGGTTCTACCACCAGAAatcagtataggcacgtttttttaaatattttttctg | *hxk-3* homology repair template; full gene deletion repair template |

**Table S2.** *C. elegans* strains used in this study.

| Strain | Genotype | Source |
|---|---|---|
| DCR9893 | hxk-2(ola589) I | This study |
| DCR9894 | hxk-3(ola590) IV | This study |
| DCR10029 | olaEx5731[flp-6p::iG6PSnFR:: mScarlet-I3; gcy-5p::mTagBFP] | This study |
| DCR10030 | hxk-2(ola589) I; hxk-3(ola590) IV; olaEx5731[flp-6p::iG6PSnFR::mScarlet-I3; gcy-5p::mTagBFP] | This study |
